# A candidate reference method for the quantification of α-synuclein in cerebrospinal fluid using an SI traceable primary calibrator and multiple reaction monitoring

**DOI:** 10.1101/2024.03.20.585804

**Authors:** Leran Zhang, Eva Illes-Toth, Adam Cryar, Giles Drinkwater, Lucia Di Vagno, Marie-Laure Pons, Julia Mateyka, Bryan McCullough, Eli Achtar, Cailean Clarkson, Laura Göschel, Peter Körtvélyessy, Chris Mussell, Christopher J. Hopley, Agnes Flöel, Christophe Hirtz, Sylvain Lehmann, Milena Quaglia

## Abstract

**Objectives:** α-synuclein aggregation is an indicator of neurodegenerative diseases such as Parkinson’s disease (PD) and recent advances have suggested that this protein could serve as a potential biomarker. It has been indicated that soluble and oligomeric α-synuclein in biological fluids could have diagnostic applications for PD. Clinical laboratories currently rely on antibody-based assays to detect α-synuclein. These assays have limited specificity, low sensitivity and poor inter-lab reproducibility, which prevents the validation of α-synuclein as a biomarkers. This study aims to fill the unmet need for the standardisation of clinical measurements for α-synuclein.

**Methods:** We report the first candidate reference method for α-synuclein, using an SI traceable primary calibrator for α-synuclein and isotope dilution mass spectrometry. The primary calibrator was traceably quantified utilising a combination of amino acid analysis and nuclear magnetic resonance. A targeted sample clean-up procedure involving a non-denaturing Lys-C digestion and solid-phase extraction allowed for the sensitive detection of multiple proteotypic α-synuclein peptides in cerebrospinal fluid (CSF) samples.

**Results:** The candidate reference method procedure showed linearity across three orders of magnitude, covering the physiological levels of α-synuclein in CSF (LOQ = 0.1 ng/g). The method was used to quantify a cohort of CSF samples and the measurements were correlated with immunoassay-based quantifications.

**Conclusions:** The SI traceable quantification of α-synuclein in complex biological matrices means that the role of this protein can be further elucidated in synucleinopathies. This candidate reference method would lead to the harmonisation of α-synuclein measurements, which may allow for development of high throughput clinical tests.

## Introduction

α-synuclein is expressed predominately in the presynaptic sites of several neurotransmitter systems in the central nervous system (1). The aggregated form of α-synuclein is the main constituent of Lewy bodies and has been implicated in both familial and sporadic forms of Parkinson’s Disease (PD) and Lewy body dementia. Thought to be one of the main pathological causes of the two diseases, α-synuclein is of interest as a potential therapeutic target and has been studied as a clinical biomarker (2). A large body of literature has also focused on α-synuclein’s potential as both diagnostic and prognostic PD biomarker and Lewy body dementia (3–6). Due to the close proximity of cerebral spinal fluid (CSF) to the brain, CSF represents one of the most tractable fluids for monitoring molecular changes occurring in the brain within a clinical setting. Recent publications have demonstrated there is a trend for lower total α-synuclein concentrations in CSF for PD patients vs healthy controls (7–10) and more recently RT-QuIC data showed the potential of α-synuclein to discriminate disease vs healthy state (11,12). Translation of these results into reliable clinical diagnostics is constrained by the variability of current antibody-based quantification methods and by the lack of analytical reference procedures to ensure spatiotemporal traceability of the results using the International System of Units (SI). Variability is especially apparent when the absolute concentrations of α-synuclein in CSF are compared. Two mass spectrometry-based methods were presented as viable alternatives to measure the absolute concentration of α-synuclein in CSF (13,14). Interestingly, despite being similar methodologies, the two publications reported an average α-synuclein concentration of 0.14 – 0.46 ng/mL (13) and 1.46 – 2.04 ng/mL (14) in measured CSF samples demonstrating that the use of mass spectrometry is not inherently a solution to overcome measurement variability. Despite the considerable progress made in measuring α-synuclein, its relevance in clinical practice as a diagnostic or prognostic marker is still controversial to date. Both immunoassay and mass spectrometry methods measure the total amount of synuclein, with the capability of mass spectrometry to differentiate between different isoforms (15). More recently, the potential of measuring misfolded states of α-synuclein showed great potential in clinical practice through the application of RT-QuIC measurements and seed amplification assays (12,16). Nevertheless, challenges in the reproducibility of calibration curves and characterisation requirements for the calibrators were observed. Therefore, there is an unmet need for an SI traceable reference method for the detection of α-synuclein in patient samples.

Analytically validated reference measurement procedures are recognised routes to the standardisation of clinical measurements and will be crucial in minimising measurement variability observed for α-synuclein in CSF. Metrological traceability, whereby quantitative measurements are directly anchored to the SI, is an established route to method specificity and inter-lab comparability. The current conventional metrological methods for the development of primary calibrators focus on the determination of the total amount of protein via amino acid analysis (AAA) and purity assessment. It is however critical that for α-synuclein, the higher order structure of the protein is considered, particularly if methods such as RT-QuIC are to be further developed into more widespread clinical applications and uptake.

Herein we report the production and application of an SI traceable primary calibrator for α-synuclein to be used for the development of a reference method. Using this primary calibrator, we have developed the first liquid chromatography-mass spectrometry candidate reference method for the quantification of α-synuclein in CSF. The method was used to quantify fifteen CSF samples and the results were compared with immunoassay data.

## Methods

### Amino acid analysis

Amino acid quantification was performed by microwave assisted hydrolysis and exact matching isotope dilution mass spectrometry (IDMS) with calibration against SI traceable amino acid standards by using a previously described method (17). SI traceability was achieved by using amino acid certified reference materials from the National Metrology Institute of Japan (NMIJ). Briefly three sample blends containing the isotopically labelled amino acids and the sample to be quantified (α-synuclein or peptide internal standards) were gravimetrically prepared together with three calibration blends containing the natural and the isotopically labelled standard amino acids in an equimolar amount to the sample blend. Reagent blanks, natural amino acid standard solution without addition of labelled standard and labelled amino acid standard solution without addition of natural standard were prepared. All sample solutions were freeze dried in a Genevac EZ-2 plus vacuum centrifuge (SP Scientific, PA, USA) and hydrolysed by acid hydrolysis in an Ethos EZ Microwave Digestion System (Analytix, Tyne and Wear, UK) with 6 M high purity hydrochloric acid. Samples were derivatized for 1 hour at 85 °C using N-tert-Butyldimethylsilyl-N-methyl-trifluroacetamide with 1% tert-Butyldimethylchlorosilane (Merck, NJ, USA) and immediately analysed by GC-MS. Analysis of the sample solutions was performed on an Agilent 5975C Mass Spectrometer (Agilent Technologies, CA, USA) using a Zebron ZB-5HT Inferno GC column (30 m x 0.25 mm i.d., 0.25 mm film thickness) with a Z- guard column (2 m, 0.53 mm i.d.) (Phenomenex, CA, USA) coupled to an Agilent 5890A gas chromatography system with CTC CombiPAL autosampler. Sample blends were bracketed with calibration blends and injected five times.

### SI traceable quantification of α-synuclein in solvent standards

Aliquots of α-synuclein (Promise Proteomics, Grenoble, France) at a nominal concentration of 0.1 µM were prepared and stored at −80 °C. Quantification was performed in triplicates by tryptic digestion and exact matching IDMS using peptides as internal standards. SI traceability was achieved through the utilisation of in-house SI traceable quantified synthetic peptides. Four peptides (MDVFMK_T1, QGVAEAAGK_T6, EGVLYVGSK_T8, TVEGAGSIAAATGFVK_T13) and four isotopically labelled analogues (all K(13C6, 15N2) labelled) were purchased from GL Biochem (Shanghai, China) with a stated purity ≥95%. Inhouse assessment of the purity of the peptides was carried out as previously described (18). Traceability was ensured by AAA and NMR. The calculated purity (as a percentage) of peptides was: MDVFMK: 95.1 ± 0.6; QGVAEAAGK: 78.7 ± 0.8; EGVLYVGSK: 99.0 ± 0.2; TVEGAGSIAAATGFVK: 26.1 ± 0.4. Additional purification of T13 was conducted, but the peptide was not used for quantification of the primary calibrator due to a different kinetic of release when compared with T1, T6 and T8. For the quantification of α-synuclein sample blends, three calibration blends and reagent blanks were gravimetrically prepared (19,20). Each sample and calibration blend was digested in duplicate for calculating the contribution of digestion variability to the measurement uncertainty. 2.4 µg of Lys-C and 4.8 µg of trypsin were added to solutions containing a nominal concentration of α-synuclein corresponding to 0.3 µg/g. Digestion was performed for 4.5 h at 37 °C with gentle agitation. Following tryptic digestion, formic acid was added to each blend to a final concentration of 1% to stop the digest. Each blend was analysed (1 μL injection) on a Waters H-Class UPLC system (Waters, Wilmslow, UK) coupled to a Waters Xevo G2-XS QToF operated in sensitivity mode with a typical resolving power of 25,000. Peptides were separated by reverse phase chromatography using a CORTECS 90 Å C18 3.0 x 150 mm column (Waters, Wilmslow, UK). Sample blends were bracketed with calibration blends and were injected five times.

### Sample preparation for the candidate reference method

CSF (0.5 mL), the α-synuclein primary calibrator (33 nM) and labelled α-synuclein (3.5 nM) were thawed at room temperature for 30 min. Nine calibration samples in 500 µL of artificial CSF (aCSF) with 0.175 mg/mL BSA were gravimetrically prepared from the primary calibrator at 0.1, 0.5, 1, 1.5, 2, 4, 6, 8 and 10 ng/g, along with the addition of labelled α-synuclein (100 µL). Tris buffer (1 M, pH 8.2) was added to a final concentration of 50 mM. Samples were digested overnight at 27 °C using Lys-C (1.17 µg). Solid phase extraction (SPE) cartridges (Phenomenex StrataX 33 µm, 30 mg/mL) were conditioned with MeOH (1 mL), MeCN (1 mL) and ammonium formate (10 mM, pH 10, 2 mL). Samples were loaded and washed with ammonium formate (10 mM, pH 10, 2 mL). Elution was performed at pH 10, using ammonium formate buffer (10 mM) with different percentages of MeCN. The T6 peptide was eluted using 3% MeCN (1.8 mL) and T8 was eluted using 10% MeCN (3 mL). The T12 and T13 peptides were eluted using 15% MeCN (1.8 mL) and combined with the T6 fraction. Samples were dried to completion overnight under vacuum at 37 °C. Samples were resuspended in 0.1% FA, 3% MeCN (70 µL), vortexed, sonicated (5 min) and centrifuged (3000 × g, 5 min). Final peptide samples were transferred into total recovery vials (Waters, Wilmslow, UK) and centrifuged (3000 × g, 5 min).

### LC-MS/MS analysis for the candidate reference method

Samples were analysed using an M-Class UPLC and a TQ-XS triple quadrupole mass spectrometer (Waters, Wilmslow, UK) in positive ionization mode. 3 × 16 µL of sample were injected on Waters HSS T3 100Å (1 × 150 mm, 1.8 µm) column at 40 °C. The mobile phase consisted of 0.1% formic acid in water (mobile phase A) and 0.1% formic acid in MeCN (mobile phase B). The gradient started at 3% mobile phase B for 2 minutes, increased to 27.6% over 17 minutes and then up to 100% over 5 minutes. Mobile phase B was pumped at 100% for 7 minutes, followed by re-equilibration at 3% for 9 minutes. Total run time for LC-MS/MS analysis was 40 min at a flow rate of 30 µL/min. MS tune parameters are as follows: Capillary voltage: 2.7 kV, Cone voltage: 40 V, Desolvation temp: 600 °C, Desolvation gas flow: 800 L/h, Cone gas flow: 250 L/h, Nebuliser: 7 bar, Source temp: 150 °C. Three multiple reaction monitoring (MRM) transitions were monitored for each peptide isotope (**Table S5**). Qualifier and quantifier ions were chosen based on their intensities and lack of isobaric interferences in the CSF pool. The ratio between the peak areas of the natural and labelled was used to create a calibration curve and to quantify the concentration of α-synuclein in the CSF samples.

## Results and Discussion

### Primary calibrator characterisation and quantification

A fundamental requirement for developing SI traceable reference methods is access to traceable value assigned standards. A commercial stock of α-synuclein (Promise Proteomics, Grenoble, France) was sourced. This stock was quantified using a tryptic digestion IDMS approach previously reported (19). Purity was confirmed to be >95% by LC-MS (**Figure S1**) Three peptides (MDVFMK: T1, QGVAEAAGK: T6 and EGVLYVGSK: T8) were selected for α-synuclein quantification based on measured digestion kinetics, with rapid enzymatic release from the standard. These peptide stocks were traceably quantified using a combination of AAA, IDMS and NMR (**Figure S2**). AAA and NMR quantifications showed good agreement with the T1 and T8 peptides. The AAA results of these two peptides were used for quantifying of the primary calibrator. However, the T6 peptide quantifications did not agree between AAA and NMR, due to the conversion of the N-terminal glutamine into pyroglutamate. The T6 NMR result was therefore used for the quantification of the primary calibrator. Following tryptic digestion optimisation and confirmation of complete enzymatic release from the protein, quantification of the purified α-synuclein stock was achieved (**Figure 1**) (21). Results gave a final α-synuclein stock concentration of 477 ng/g with a combined uncertainty (k=2) of 9.1% (**Table S1**). The combined uncertainty calculation includes uncertainties from AAA/NMR quantification, proteolytic digestion and homogeneity of the sample.

**Figure 1.**
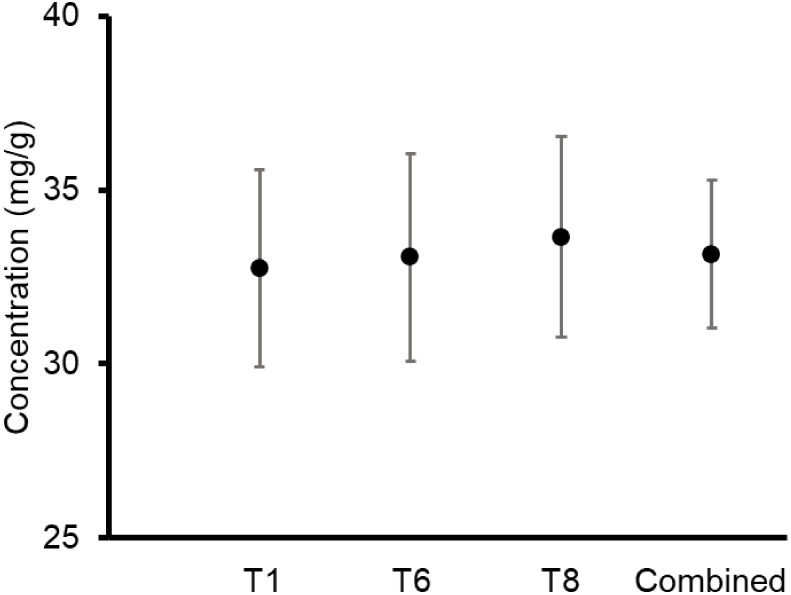
Plot showing the agreement between individual peptide level values for T1, T6 and T8 tryptic peptides liberated from the α-synuclein primary calibrator stock (n = 9). Combining the peptide level data permits value assignment of the primary calibrator with its expanded uncertainty (k = 2), representing the 95% confidence interval.

### Structural characterisation of the primary calibrator

For the reference method, the aggregation properties and structure of the primary calibrator must be characterised. Bovine serum albumin (BSA) was used as a carrier protein to reduce sample loss of the primary calibrator in the calibration curve of the candidate reference method. Therefore, structural characterisation was performed with and without BSA using ion mobility mass spectrometry under native-like conditions (**Figure S3**). A previous report suggested that human serum albumin targets α-synuclein and redirects its aggregation into different intermediates (22). No apparent changes were detected in the charge state distribution of α-synuclein encompassing ions from 6+ to 16+ suggesting that BSA can be used under our conditions. Analyses of the corresponding drift times revealed that BSA caused only a minor compaction of ions ranging from 8+ to 11+. The full series from charge states 6+ to 16 + can be found in **Figure S4**. The shifts in drift times in the presence of bovine serum albumin did not result in significant changes in the calculated mean collision cross sections (^TW^CCS_N2→He_) (**Table S2**). To probe the stability of α-synuclein with and without BSA, collision induced unfolding (CIU) was employed under native-like conditions. The samples showed no major differences in their CIU fingerprints (**Figure S5**). α-synuclein is known to self-assemble to oligomers which are often observed in recombinant sources (23). On closer inspection of the mass spectra, we detected various preformed oligomeric populations (**Tables S3 – S4** and **Figure S6**) of α-synuclein with very low intensities. Importantly, we did not find differences in the distributions of these different multimeric α-synuclein complexes ranging from dimers to hexamers in the absence or presence of bovine serum albumin (**Figures S7 – S9**). In order to address the effect of freeze and thaw cycles during sample preparation of the primary calibrator, we subjected 1 µM α-synuclein with 0.175 mg/mL BSA (2.6 µM) to one or two cycles of freeze and thaw. It was observed that up to two freeze thaw cycles led to no discernible differences in the charge state distributions (**Figure S10**). Lastly, brief vortexing steps were included during the preparation of the primary calibrator. To investigate whether any aggregation occurred and to study the effects of shaking on α-synuclein, 45 µL of 15 μM protein was incubated at pH 7.0 at 21 – 22 °C in the presence of 0.175 mg/mL bovine serum albumin (2.6 µM) with or without shaking at 350 rpm on an orbital benchtop shaker. Between the two conditions monitored, we observe neither significant changes, nor any indications of aggregation, which often exhibits bimodal distributions as judged by intact hydrogen deuterium exchange mass spectrometry (**Figure S11**).

### Quantification of α-synuclein in CSF

Based on the results obtained from the quantification of the primary calibrator, the T6, T8, T12 and T13 peptides were chosen as measurands for the reference method due to their sensitivity, selectivity, and equimolar release between the natural and labelled proteins (**Figure 2A**). Previous literature reports CSF α-synuclein concentrations to be in the region of low ng/mL, placing the required method limit of quantification (LOQ) at the sub ng/mL (14). Mass spectrometry is a highly sensitive analytical technique, where attomole concentrations can be detected in solvent standards. At low concentrations in complex biological samples however, an optimised sample preparation protocol is required to reduce ion suppression and the presence of closely eluting isobaric compounds which may introduce bias. To meet these requirements, a bespoke sample clean up method was developed (**Figure 2B**). Initial experiments confirmed the previously reported use of a non-denaturing and non-reducing overnight tryptic digestion protocol for increased method sensitivity towards α-synuclein in CSF. A high pH SPE protocol was optimised to allow the concurrent removal of intact protein, salts and other small molecules. The monitored peptides (T6, T8, T12 and T13) were fractionated during SPE elution using increasing concentrations of acetonitrile. The T6, T12 and T13 peptides were collected in the same fraction because T6 elutes early in the chromatographic gradient, while T12 and T13 co-elutes near the end of the gradient. This significantly reduces analysis time. Samples were concentrated and injected in triplicate into an UPLC system with a forty-minute acidic reverse phase gradient prior to MRM MS analysis. Both standard flow and capillary flow chromatographic methods were developed and compared. The capillary method produced higher signal to noise (S/N) and was taken forward for sample analysis. MRM transitions with the best signal intensity along with their collision energies were optimised using Skyline (24) (**Table S5**).

**Figure 2.**
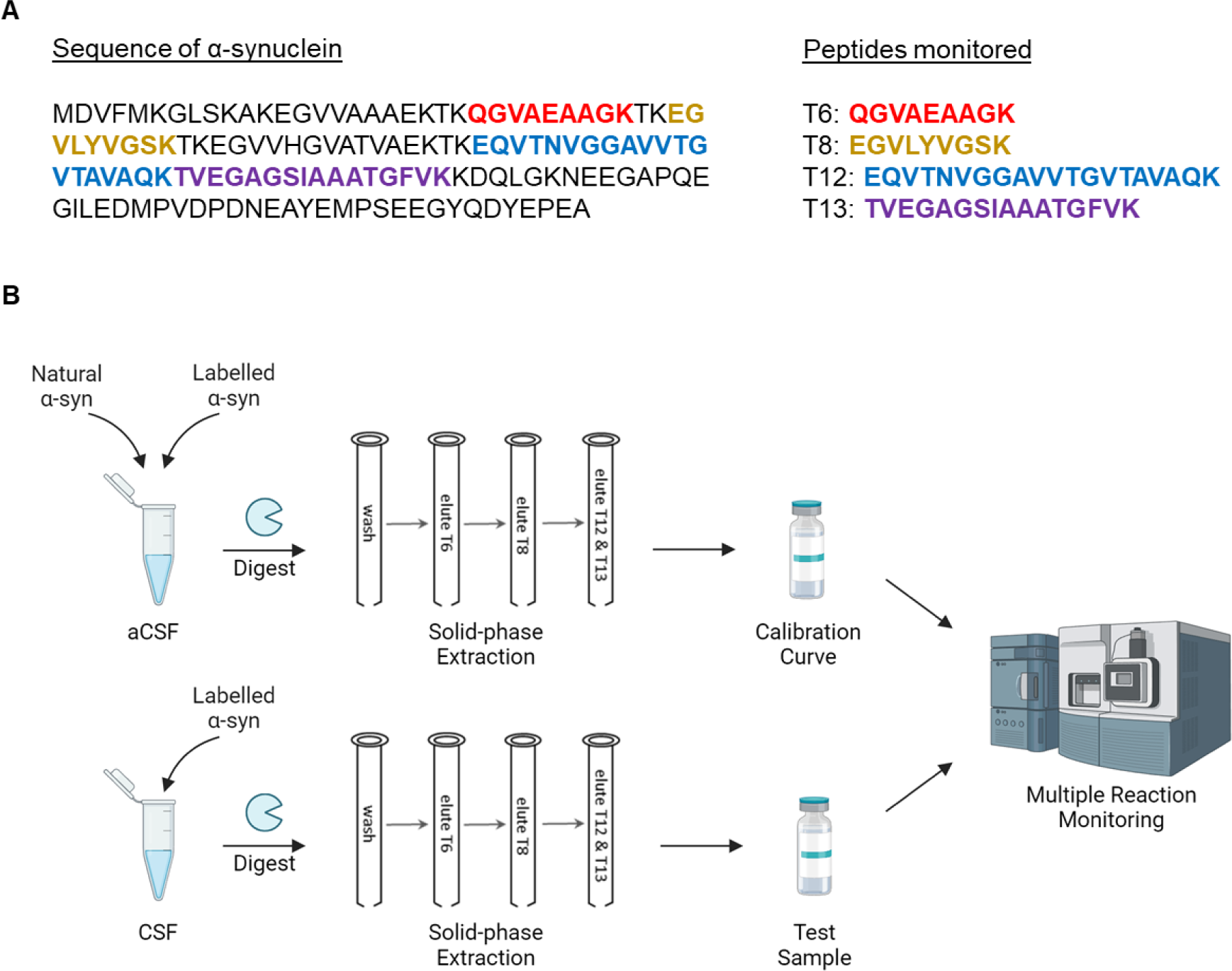
Protocol for the candidate reference method procedure. (A) Sequence of α-synuclein and the monitored peptides. (B) Sample preparation and analysis workflow for artificial CSF (aCSF) or real CSF.

Initial trials applying the above method showed no quantifiable signal with the T6, T12 and T13 peptides in CSF. It was hypothesised that despite the non-reducing tryptic digestion, matrix suppression was still an issue with these peptides. To address this, a more specific protease was required for the digestion of α-synuclein to reduce the complexity of the sample. α-synuclein is unique in the fact that it does not contain any arginine residues. Therefore, the use of Lys-C allowed for the full digestion of α-synuclein while background proteins would only be partially digested compared to a tryptic digestion. Furthermore, Lys-C has a greater digestion efficiency for lysine, which would in theory allow for a more complete digestion of the measurand.

The switch from trypsin to Lys-C resulted in a four to thirty-fold increase in sensitivity compared to trypsin across the four peptides in spiked samples, with T12 and T13 showing the best improvements. In real CSF samples, matrix effects of T12 and T13 were reduced significantly to a point where the peak intensities were comparable to aCSF. Unfortunately, the T6 peptide was still poorly detectable in CSF. The optimised method showed excellent linearity across three orders of magnitude (0.1 ng/g – 10 ng/g) of α-synuclein, with r^2^ = 0.999 for all four peptides (**Figure 3A**). The LOQ was assigned to be the lowest point on the calibration curve (0.1 ng/g), which fulfils the requirement of having a sub ng/mL method. Precision between aCSF injections was between 1.1% – 5.8% (**Figure 3B**). Intra-assay precision between six replicates of the same CSF sample were satisfactory for the T8, T12 and T13 peptides (2.8% – 6.4%). CSF samples that have been over spiked with primary calibrator to 5 ng/g showed even higher precision (2.1% – 2.2%). However, T6 peptides displayed poor sensitivity and was excluded in follow up experiments. Measurement uncertainty calculations are presented in the supporting information.

**Figure 3.**
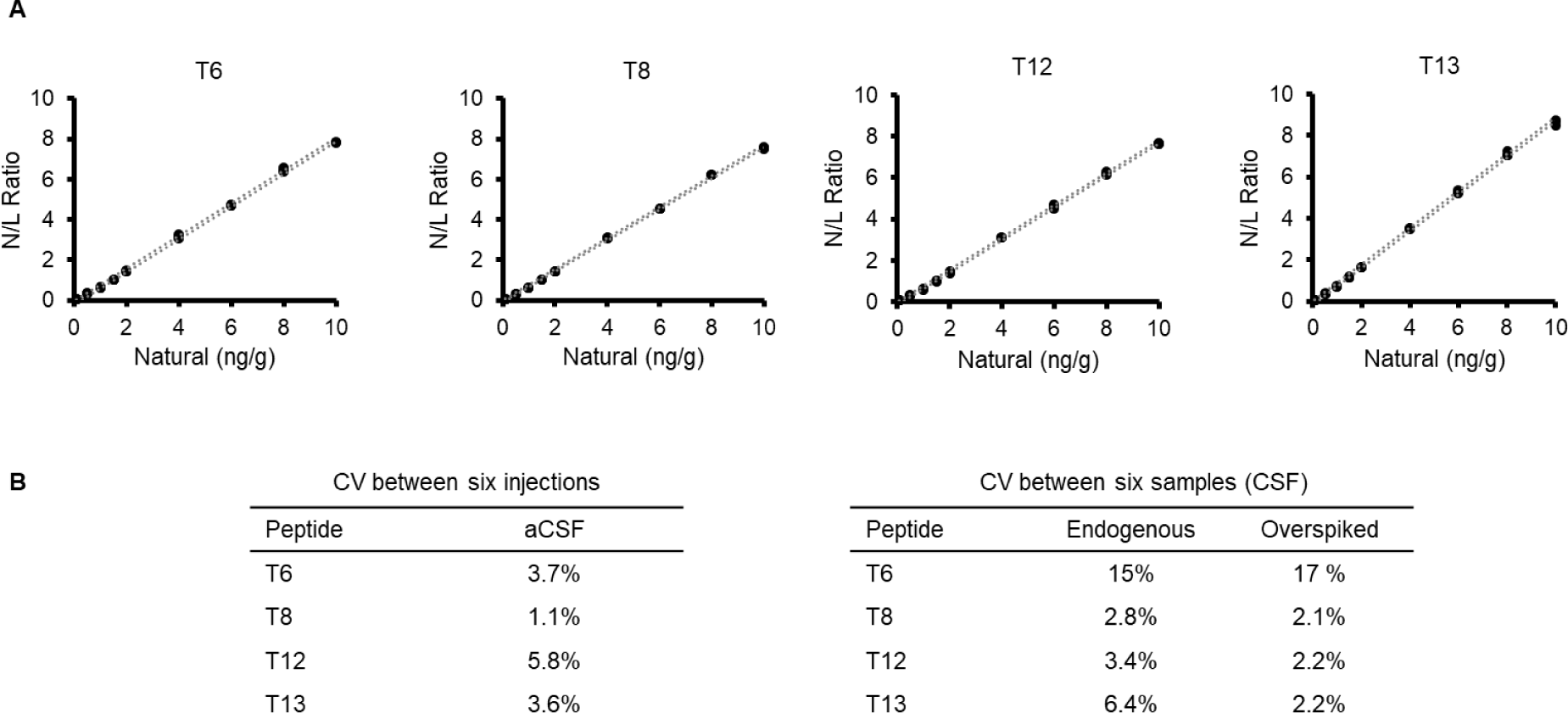
Validation experiments for the candidate reference method. (A) Calibration curves for the four monitored peptides (T6, T8, T12 and T13), plotting the natural/labelled peak area ratio against the volumetrically prepared natural concentration. Upper and lower limits of uncertainty at 95% confidence are represented by dotted lines. r^2^ = 0.999 for all peptides. (B) Method precision between injections in aCSF and between samples in real CSF.

### Quantification of α-synuclein in fifteen CSF samples

Fifteen CSF samples from Charité Universitätsmedizin Berlin were analysed using the candidate reference method procedure (**Figure 4A**). These samples were first analysed via immunoassay. 500 µL of each sample was used for MS analysis. Calibration curves showed good linearity for all four peptides (r^2^ = 0.996 – 0.999). The full calibration curve covered three orders of magnitude (0.1 ng/g – 10 ng/g), but only the lower half was used for the quantification of α-synuclein in the CSF samples. The curves were plotted using the gravimetrically calculated concentration of primary calibrator against the ratio of peak areas between the labelled and natural MRM transitions. The amount of α-synuclein in the fifteen samples were calculated using the same ratios. The method monitored three peptides, T8 (which is a sequence that is shared with ß-synuclein), T12 and T13 (**Figure 4B**). Although, the T6 calibration curve is in line with the other peptides, the T6 peptide was not detected in the CSF samples. This may be due to the CSF matrix suppressing the peptide signal.

**Figure 4.**
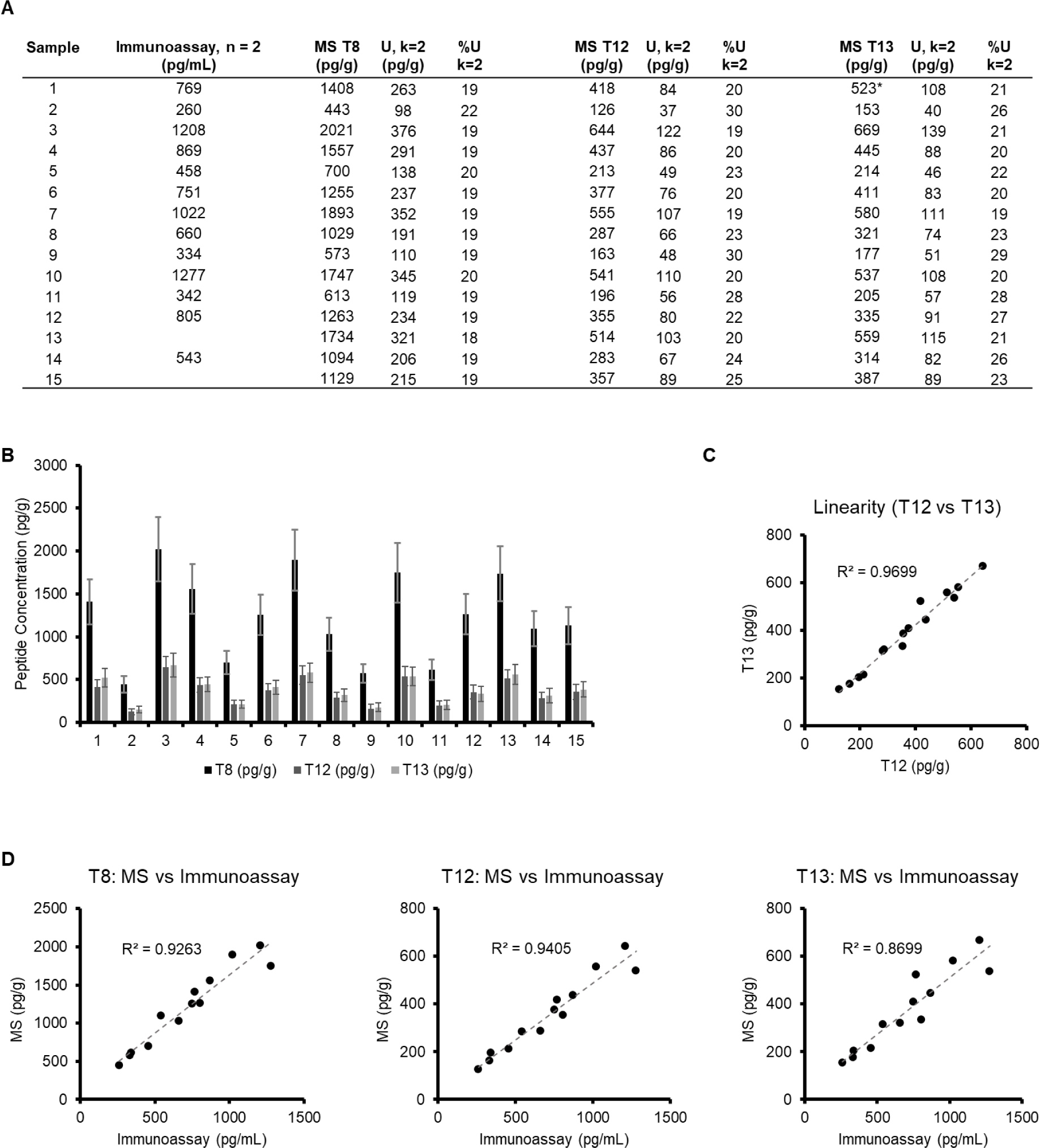
Analysis of a panel of CSF samples from Charité. (A) Levels of α-synuclein in the CSF samples analysed from an immunoassay and the mass spectrometry method (MS). * = omitted result due to isobaric interferences. (B) Comparison of concentration values obtained from the T8, T12 and T13 peptides. Error bars represent the expanded uncertainty at 95% confidence. (C) T12 and T13 values were plotted to show linearity. (D) Scatter plot comparing the quantification results from the immunoassay and the MS method for the T8, T12 and T13 peptides.

The calculated concentration values between T12 and T13 showed good linearity (r^2^ = 0.97), with Sample 1 being the only outlier (**Figure 4C**). A thorough inspection of the peak areas of this sample showed that the natural channel had a slightly higher peak area compared to the other two MRMs of the peptide, suggesting the presence of an isobaric interference. Further analysis of the ratios between the main and qualifier MRMs of the other CSF samples have shown that only the aforementioned sample had significant isobaric interferences (**Table S6**).

The values associated with the immunoassay were consistently above the values obtained from the T12 and T13 peptides, while being below the T8 values obtained by the MS method. This suggests that the immunoassay may not be fully specific and may be detecting other proteins such as ß-synuclein. Furthermore, since the monoclonal antibodies are specific to two epitopes (Glu110-Tyr125 and Val15-Tyr125 chains), it is limited to detecting certain folding or aggregation states of α-synuclein, while the mass-spectrometry method is less affected by protein structure. This may explain the lower quantification values for the T12 and T13 peptides in the immunoassay. A degree of linearity was observed for all three peptides when plotting the immunoassay results with each of the individual peptides (**Figure 4D**).

## Conclusion

We report an SI traceable candidate reference method for the quantification of α-synuclein from CSF samples. The method covers the physiological concentrations of α-synuclein over three orders of magnitude (0.1 ng/g – 10 ng/g). The candidate reference method showed correlation with immunoassay results while analysing fifteen patient samples. Furthermore, the structurally characterised SI traceable primary calibrator of α-synuclein can be applied in other applications such as in phenotypic assays or for the advancement of RT-QuIC into the clinic. Although the reference method requires further validation in terms of recovery, linearity and precision, it serves as a proof-of-concept that α-synuclein can be traceably measured in complex biological systems.

## Acknowledgements

Funding was provided by the EMPIR programme co-financed by the Participating States and from the European Union’s Horizon 2020 research and innovation programme (15HLT09-NeuroMET and 18HLT09-NeuroMET2). Samples for generating CSF pools were retrospectively selected from the NeuroCognition Biobank of Montpellier’s University Hospital from patients recruited in the Resource and Research Memory Center. Patients gave informed and written consent to have their samples stored in an officially registered and ethically approved biological collection (#DC-2008-417) and later used for scientific research. For the CSF panel, all participants gave written informed consent before participation in the NeuroMET and NeuroMET2 studies. The study was approved by the ethics committee of the Charité university hospital (EA1/197/16 and EA2/121/19).

## Abbreviations

AAA: Amino acid analysis
aCSF: Artificial cerebrospinal fluid
CCS: Collision cross section
CIU: Collision induced unfolding
CNS: Central nervous system
CSF: Cerebrospinal fluid
EDTA: Ethylenediaminetetraacetic acid
FA: Formic acid
GC-MS: Gas chromatography mass spectrometry
HPLC: High performance liquid chromatography
IDMS: Isotope dilution mass spectrometry
IE: Ion exchange
IPTG: Isopropyl β-D-1-thiogalactopyranoside
LC-MS: Liquid chromatography mass spectrometry
LOQ: Limit of quantification
MeCN: Acetonitrile
MRM: Multiple reaction monitoring
MS: Mass spectrometry
NMIJ: National Metrology Institute of Japan
NMR: Nuclear magnetic resonance
PD: Parkinson’s disease
RT-QuIC: Real-time quaking-induced conversion
SEC: Size exclusion chromatography
SI: International System of Units
SPE: Solid phase extraction
UPLC: Ultra-performance liquid chromatography

