## Supporting Information for "A candidate reference method for the quantification of α-synuclein in cerebrospinal fluid using an SI traceable primary calibrator and multiple reaction monitoring"

(1) LGC Group, Queens Road, TW11 0LY Teddington, UK.
(2) IRMB-PPC, INM, CHU Montpellier, INSERM, CNRS, Université de Montpellier, 34295 Montpellier, France
(3) Charité - Universitätsmedizin Berlin, Corporate Member of Freie Universität Berlin and Humboldt-Universität zu Berlin, Department of Neurology, 10117 Berlin, Germany,
(4) Charité - Universitätsmedizin Berlin, Corporate Member of Freie Universität Berlin and Humboldt-Universität zu Berlin, Neuroscience Clinical Research Center, 10117 Berlin, Germany
(5) Labor Berlin, Innovations, Sylter Strasse 2, 13353 Berlin, Germany
(6) Department of Neurology, University Medicine Greifswald, 17475 Greifswald, Germany

Table of contents

**Figure S1**. Purity of the primary calibrator
**Figure S2.** Quantification of the T1, T6 and T8 peptide stocks
**Table S1.** SI traceable quantification values for the primary calibrator
**Figure S3.** Mass spectrum of α-syn without or with BSA under native-like conditions
**Figure S4.** Drift time distributions of α-syn without or with BSA
**Table S2**. Mean $^{TW}CCS_{N2\rightarrow He}$ ± one STD of α-syn without or with BSA
**Figure S5.** CIU plots of α-syn without or with BSA
**Table S3**. Different theoretical monomeric and oligomeric masses (Da) of α-syn
**Figure S6.** Native top-down CID fragmentation mass spectrum of α-syn (m/z 2893 ion)
**Figure S7**. Expanded mass spectrum of 1 μM α-syn
**Figure S8**. Expanded mass spectrum of 1 μM α-syn with BSA
**Figure S9**. Deconvoluted mass of α-syn
**Figure S10.** Mass spectrum of α-syn with BSA following freeze and thaw

**Figure S11.** Deuterium uptake of α-syn in the presence of bovine serum albumin with or without shaking
**Table S5**. Skyline MRM optimisation
**Table S6.** Isobaric interferences in CSF samples
**Table S7.** Recovery values (%) of four α-syn synuclein peptides.
**S12.** Materials and Data Processing

**Supplementary figures**

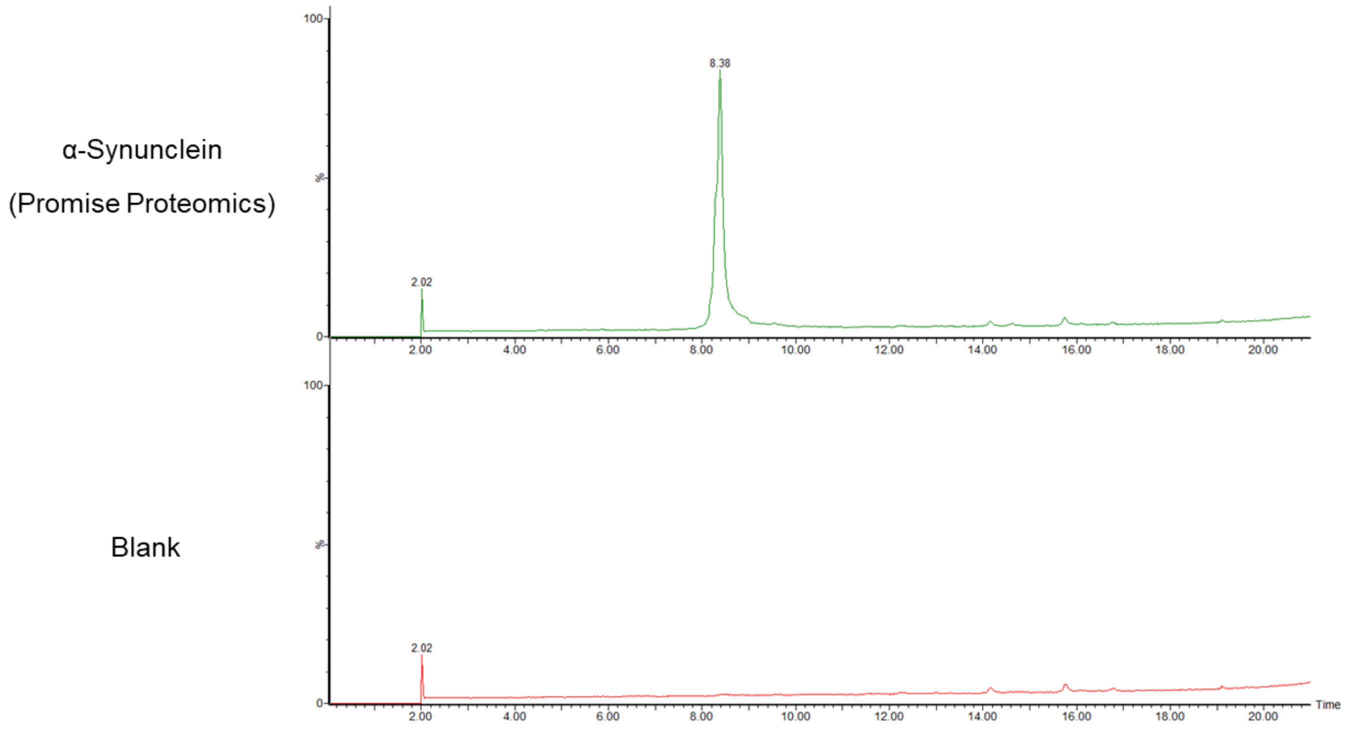


**Figure S1.** Purity of the primary calibrator. Chromatograms of a blank and the primary calibrator. Samples were analysed using an H-Class UPLC (Waters) and a Xevo G2-XS qToF mass spectrometer (Waters) in positive ionisation mode the mobile phase consisted of 0.5% formic acid in water (mobile phase A) and 0.5 % formic acid in MeCN (mobile phase B). The gradient starts from 15% mobile phase B for 1 minute, increased to 80% over 19 minutes, and then up to 95% over 1 minute. Mobile phase B was pumped at 95% for 3 minutes, followed by gradient to 15% over 1 minute. The column was re-equilibrated at 15% mobile phase B for 10 minutes. Total run time was 35 minutes at a flow rate of 0.25 ml/min.

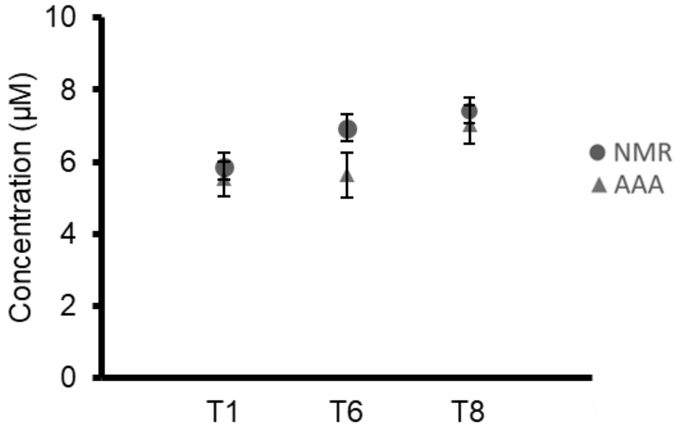


**Figure S2.** Quantification of the T1, T6 and T8 peptide stocks. Peptide stocks were quantified using both amino acid analysis (AAA) or nuclear magnetic resonance (NMR). Error bars indicate two standard deviations of replicate analyses.

### Quantification of the primary calibrator

**Table S1.** SI traceable quantification values for the primary calibrator (mg/g) from three sample blends, injected three times. The uncertainty (u) and percentage uncertainty (%u) are also presented.

| Blend | T1 | u | %u | T6 | u | %u | T8 | u | %u | Av | u | %u |
|---|---|---|---|---|---|---|---|---|---|---|---|---|
| 1 | 35.52 | 0.48 | 1.36% | 36.00 | 0.54 | 1.51% | 36.77 | 0.31 | 0.85% | 36.10 | 0.79 | 2.19% |
| 1 | 32.72 | 0.39 | 1.18% | 33.29 | 0.63 | 1.90% | 33.97 | 0.32 | 0.96% | 33.33 | 0.81 | 2.43% |
| 1 | 31.77 | 0.46 | 1.44% | 32.38 | 0.55 | 1.68% | 33.04 | 0.32 | 0.96% | 32.40 | 0.78 | 2.41% |
| 2 | 33.56 | 0.41 | 1.21% | 33.49 | 0.61 | 1.84% | 34.35 | 0.40 | 1.16% | 33.80 | 0.84 | 2.48% |
| 2 | 31.96 | 0.34 | 1.06% | 32.26 | 0.61 | 1.90% | 32.90 | 0.42 | 1.29% | 32.37 | 0.82 | 2.53% |
| 2 | 28.60 | 0.35 | 1.24% | 28.76 | 0.74 | 2.56% | 29.24 | 0.33 | 1.13% | 28.87 | 0.88 | 3.06% |
| 3 | 38.17 | 0.76 | 2.00% | 38.95 | 0.88 | 2.25% | 38.90 | 0.43 | 1.11% | 38.67 | 1.24 | 3.21% |
| 3 | 30.02 | 0.41 | 1.35% | 30.41 | 0.68 | 2.24% | 30.93 | 0.46 | 1.49% | 30.45 | 0.92 | 3.01% |
| 3 | 32.47 | 0.47 | 1.45% | 32.09 | 0.53 | 1.67% | 32.71 | 0.36 | 1.10% | 32.42 | 0.80 | 2.46% |

### Structural characterisation of the primary calibrator

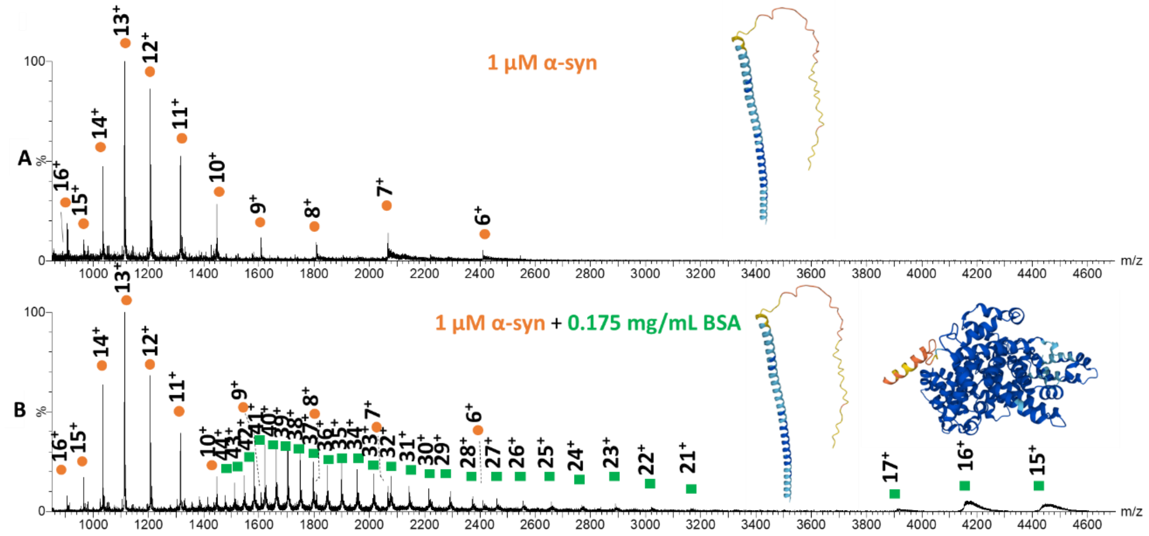


**Figure S3**. A: Mass spectrum of 1 µM α-syn in 100 mM ammonium acetate, pH 7.0 under native-like conditions (depiction of α-syn from Alphafold: AF-P37840-F1). B: Mass spectrum of 1 µM α-syn with 0.175 mg/mL BSA (2.6 µM) in 100 mM ammonium acetate (depiction of BSA from Alphafold: AF-P02769-F1), pH 7.0 under native-like conditions. α-syn charge states are shown with orange circles, bovine serum albumin monomers with green squares.

This broad charge state distribution with a multimodal appearance is typical of an intrinsically disordered protein with large structural flexibility(1–3).

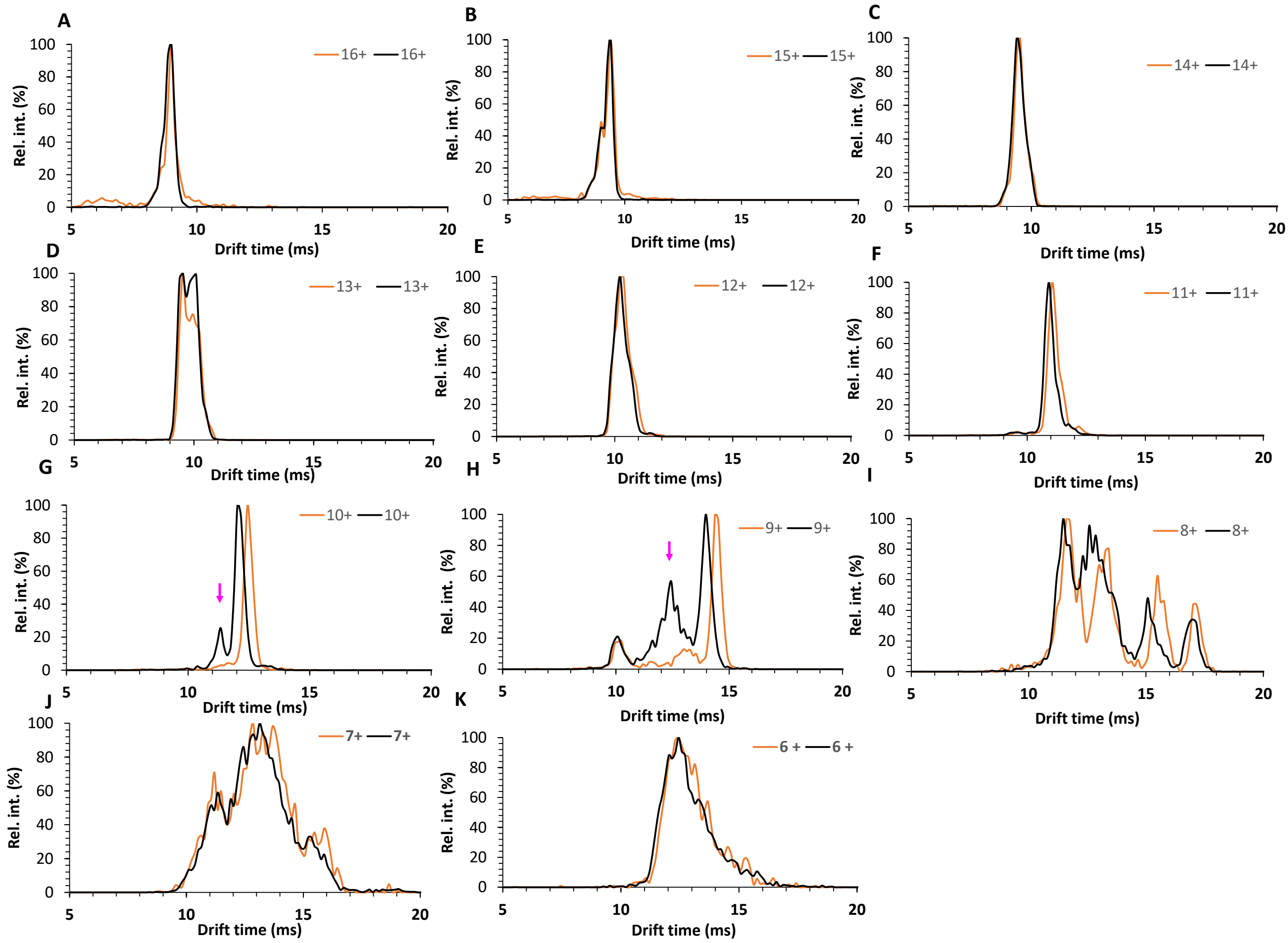


**Figure S4.** A, Drift time distributions of A, 16+ of 1 μM α-syn only (orange) and that in the presence of bovine serum albumin (black); B, 15+ of 1 μM α-syn only (orange) and that in the presence of bovine serum albumin (black); C, 14+ of 1 μM α-syn only (orange) and that in the presence of bovine serum albumin (black); D, 13+ of 1 μM α-syn only (orange) and that in the presence of bovine serum albumin (black); E, 12+ of 1 μM α-syn only (orange) and that in the presence of bovine serum albumin (black); F, 11+ of 1 μM α-syn only (orange) and that in the presence of bovine serum albumin (black); G, 10+ of 1 μM α-syn only (orange) and that in the presence of bovine serum albumin (black) with the arrow indicating a new conformer; H, 9+ of 1 μM α-syn only (orange) and that in the presence of bovine serum albumin (black) with an arrow indicating an increase in the abundance of intermediate conformation; I, 8+ of 1 μM α-syn only (orange) and that in the presence of bovine serum albumin (black); J, 7+ of 1 μM α-syn only (orange) and that in the presence of bovine serum albumin (black); K, 6+ of 1 μM α-syn only (orange) and that in the presence of bovine serum albumin (black); All data shown were acquired at 38 V IMS wave height.

Note the appearance of a new, minor conformer with a drift time of 11.34 ms that becomes visible and distinguishable in addition to the most dominant conformer of 10+ ion with 12.02 ms. Of note, the relative distribution of 9+ conformers was also different in the presence of bovine serum albumin with an increase in the intermediate conformer relative to the most dominant conformer. All $^{TW}CCS_{N2\rightarrow He}$ can be found in **Table S2**.

**Table S2**. Mean $^{TW}CCS_{N2\rightarrow He}$ ± one STD of α-syn without or with BSA in 100 mM $NH_4OAc$ at pH 7, n=3 and reference CCS. Data were acquired on a Synapt G2 Si mass spectrometer in positive ionisation, in sensitivity mode. Literature values measured in negative mode denoted with a superscript (-).

| ID | 1 µM α-syn 100 mM $NH_4OAc$ | | 1 µM α-syn 100 mM $NH_4OAc$ + 2.6 µM bovine serum albumin | | |
|---|---|---|---|---|---|
| z | $^{TW}CCS_{N2\rightarrow He}$ (Å$^2$) | ± STD (Å$^2$) | $^{TW}CCS_{N2\rightarrow He}$ (Å$^2$) | ± STD (Å$^2$) | Literature CCS (Å$^2$) |
| 20 | - | - | - | - | **2620**(4) |
| 19 | - | - | - | - | **2605 ± 183**(4)<br>**~3340**(5) |
| 18 | - | - | - | - | **2742 ± 11**(4)<br>**~3900**(6)<br>**~3350**(5) |
| 17 | - | - | - | - | **2617 ± 153**(4)<br>**~3750** (6)<br>**~3200**(5) |
| 16 | **3083** | **12** | **3083** | **12** | **2560 ± 229**(4)<br>**~3600**(6)<br>**~3030**(5) |
| 15 | **2999** | **8** | **2999** | **8** | **2476 ± 161**(4)<br>**~3400**(6)<br>**~2940**(5) |
| 14 | **2832** | **5** | **2821** | **19** | **2446 ± 136**(4)<br>**~3200**(6)<br>**~2650**(5) |
| 13 | **2629** | **5** | **2629** | **5** | **~3000**(7)<br>**2150 ± 398**(4)<br>**~2900**(6)<br>**~2650**(5) |
| | - | - | - | - | **~2680**(5) |
| | **2733** | **24** | **2742** | **20** | **~3250**(6)<br>**~2750**(5) |
| 12 | **2579** | **9** | **2570** | **8** | **2311 ± 234**(4)<br>**~3000**(6)<br>**~2600**(5) |
| | - | - | - | - | **~2750**(5) |
| 11 | **2492** | **1** | **2484** | **13** | **2161 ± 173**(4)<br>**~2750**$^{(-)}$(1)<br>**~2875**(6)<br>**~2480**(5) |
| | - | - | - | - | **~2640**(5) |
| 10 | **2487** | **7** | **2450** | **32** | **~1800**(7)<br>**1951 ± 155**(4)<br>**~2500**$^{(-)}$<br>**~2850**(6)<br>**~2450**(5) |
| | - | - | **2327** | **14** | - |
| 9 | **1897** | **14** | **1897** | **14** | **1506 ± 248**(4) |
| | **2319** | **7** | **2280** | **43** | **~2400**$^{(-)}$(1)<br>**~2750**(6) |
| | **2518** | **16** | **2479** | **44** | **~2900**(6)<br>**~2460**(5) |
| 8 | **1897** | **12** | **1886** | **22** | **1333 ± 181**(4)<br>**~1470**$^{(-)}$(1)<br>**~2200** (6)<br>**~1880**(5) |
| | **2090** | **7** | **2062** | **33** | **~1550**$^{(-)}$(1) |

| | | | | | |
|---|---|---|---|---|---|
| | | | | | **~2350**(6)<br>**~2070**(5) |
| | **2371** | **18** | **2337** | **36** | **~2750**(6)<br>**~2340**(5) |
| | **2558** | **27** | **2558** | **31** | **~3000**(6)<br>**~2500**(5) |
| 7 | **1601** | **18** | **1601** | **17** | **1252 ± 147**(4)<br>**1425**$^{(-)}$(1)<br>**~1750**(6)<br>**1578**(5) |
| | **1871** | **30** | **1836** | **60** | **~1450**$^{(-)}$(1)<br>**~2250**(6)<br>**1823**(5) |
| | **2099** | **5** | **2050** | **38** | **~2500**(6)<br>**1998**(5) |
| 6 | **1504** | **15** | **1495** | **18** | **~1400**(7)<br>**1217 ± 147**(4)<br>**~1360**$^{(-)}$(1)<br>**~1760**(6)<br>**~1470**(5) |
| | - | - | - | - | **~1400**$^{(-)}$(1)<br>**~2000**(6)<br>**~1750**(5) |
| 5 | - | - | - | - | **1043 ± 133**(4)<br>**~1750**(6) |

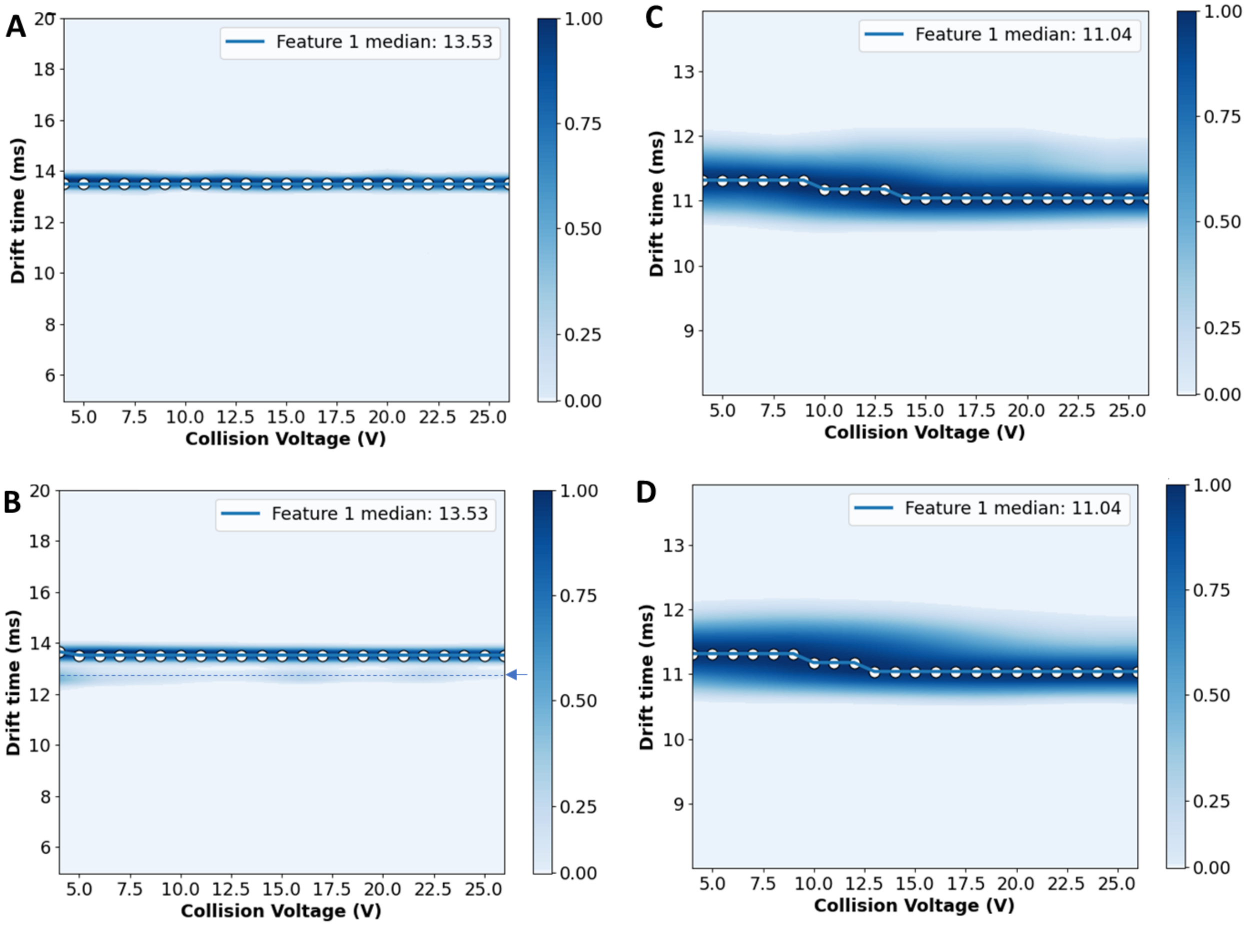


**Figure S5.** CIU plots of A, 1 μM α-syn (10+) B, 1 μM α-syn + 2.6 μM BSA (10+). The blue arrow on the right indicates an additional conformer with a more compact conformation. C, 1 μM α-syn (12+) and D, 1 μM α-syn + 0.175 mg/mL SA (12+) generated in CIU Suite 2 with a smoothing window of 5, iteration 2, and axis scaling step of 2 of collision voltage for interpolation. Voltages were ramped by 2 V increments in the trap region between 4 V- 26 V.

Similarly to a previous report(8), we did not observe notable structural transitions of α-syn by CIU.

**Table S3**. Different theoretical monomeric and oligomeric masses (Da), respective m/z and z values.

| MONOMER | 14460 | DIMER | 28920 | TRIMER | 43380 | TETRAMER | 57840 | PENTAMER | 72300 | HEXAMER | 86760 |
|---|---|---|---|---|---|---|---|---|---|---|---|
| m/z | z | m/z | z | m/z | z | m/z | z | m/z | z | m/z | z |
| 804.3 | 18 | 804.3 | 36 | 1206.0 | 36 | 1258.4 | 46 | 1206.0 | 60 | 1206.0 | 72 |
| 851.6 | 17 | 827.3 | 35 | 1240.4 | 35 | 1286.3 | 45 | 1226.4 | 59 | 1223.0 | 71 |
| 904.8 | 16 | 851.6 | 34 | 1276.9 | 34 | 1315.6 | 44 | 1247.6 | 58 | 1240.4 | 70 |
| 965.0 | 15 | 877.4 | 33 | 1315.6 | 33 | 1346.1 | 43 | 1269.4 | 57 | 1258.4 | 69 |
| 1033.9 | 14 | 904.8 | 32 | 1356.6 | 32 | 1378.2 | 42 | 1292.1 | 56 | 1276.9 | 68 |
| 1113.3 | 13 | 933.9 | 31 | 1400.4 | 31 | 1411.7 | 41 | 1315.6 | 55 | 1295.9 | 67 |
| 1206.0 | 12 | 965.0 | 30 | 1447.0 | 30 | 1447.0 | 40 | 1339.9 | 54 | 1315.6 | 66 |
| 1315.6 | 11 | 998.2 | 29 | 1496.9 | 29 | 1484.1 | 39 | 1365.2 | 53 | 1335.8 | 65 |
| 1447.0 | 10 | 1033.9 | 28 | 1550.3 | 28 | 1523.1 | 38 | 1391.4 | 52 | 1356.6 | 64 |
| 1607.7 | 9 | 1072.1 | 27 | 1607.7 | 27 | 1564.3 | 37 | 1418.7 | 51 | 1378.2 | 63 |
| 1808.5 | 8 | 1113.3 | 26 | 1669.5 | 26 | 1607.7 | 36 | 1447.0 | 50 | 1400.4 | 62 |
| 2066.7 | 7 | 1157.8 | 25 | 1736.2 | 25 | 1653.6 | 35 | 1476.5 | 49 | 1423.3 | 61 |
| 2411.0 | 6 | 1206.0 | 24 | 1808.5 | 24 | 1702.2 | 34 | 1507.3 | 48 | 1447.0 | 60 |
| 2893.0 | 5 | 1258.4 | 23 | 1887.1 | 23 | 1753.7 | 33 | 1539.3 | 47 | 1471.5 | 59 |
| | | 1315.6 | 22 | 1972.8 | 22 | 1808.5 | 32 | 1572.7 | 46 | 1496.9 | 58 |
| | | 1378.2 | 21 | 2066.7 | 21 | 1866.8 | 31 | 1607.7 | 45 | 1523.1 | 57 |
| | | 1447.0 | 20 | 2170.0 | 20 | 1929.0 | 30 | 1644.2 | 44 | 1550.3 | 56 |
| | | 1523.1 | 19 | 2284.2 | 19 | 1995.5 | 29 | 1682.4 | 43 | 1578.5 | 55 |
| | | 1607.7 | 18 | 2411.0 | 18 | 2066.7 | 28 | 1722.4 | 42 | 1607.7 | 54 |
| | | 1702.2 | 17 | 2552.8 | 17 | 2143.2 | 27 | 1764.4 | 41 | 1638.0 | 53 |
| | | 1808.5 | 16 | 2712.3 | 16 | 2225.6 | 26 | 1808.5 | 40 | 1669.5 | 52 |
| | | 1929.0 | 15 | 2893.0 | 15 | 2314.6 | 25 | 1854.9 | 39 | 1702.2 | 51 |
| | | 2066.7 | 14 | 3099.6 | 14 | 2411.0 | 24 | 1903.6 | 38 | 1736.2 | 50 |
| | | 2225.6 | 13 | 3337.9 | 13 | 2515.8 | 23 | 1955.1 | 37 | 1771.6 | 49 |
| | | 2411.0 | 12 | 3616.0 | 12 | 2630.1 | 22 | 2009.3 | 36 | 1808.5 | 48 |
| | | 2630.1 | 11 | 3944.6 | 11 | 2755.3 | 21 | 2066.7 | 35 | 1847.0 | 47 |
| | | 2893.0 | 10 | 4339.0 | 10 | 2893.0 | 20 | 2127.5 | 34 | 1887.1 | 46 |
| | | 3214.3 | 9 | 4821.0 | 9 | 3045.2 | 19 | 2191.9 | 33 | 1929.0 | 45 |
| | | | | | | 3214.3 | 18 | 2260.4 | 32 | 1972.8 | 44 |
| | | | | | | 3403.4 | 17 | 2333.3 | 31 | 2018.7 | 43 |
| | | | | | | 3616.0 | 16 | 2411.0 | 30 | 2066.7 | 42 |
| | | | | | | 3857.0 | 15 | 2494.1 | 29 | 2117.1 | 41 |
| | | | | | | 4132.4 | 14 | 2583.2 | 28 | 2170.0 | 40 |
| | | | | | | 4450.2 | 13 | 2678.8 | 27 | 2225.6 | 39 |
| | | | | | | 4821.0 | 12 | 2781.8 | 26 | 2284.2 | 38 |
| | | | | | | 5259.2 | 11 | 2893.0 | 25 | 2345.9 | 37 |
| | | | | | | 5785.0 | 10 | 3013.5 | 24 | 2411.0 | 36 |
| | | | | | | | | 3144.5 | 23 | 2479.9 | 35 |
| | | | | | | | | 3287.4 | 22 | 2552.8 | 34 |
| | | | | | | | | 3443.9 | 21 | 2630.1 | 33 |
| | | | | | | | | 3616.0 | 20 | 2712.3 | 32 |
| | | | | | | | | 3806.3 | 19 | 2799.7 | 31 |
| | | | | | | | | 4017.7 | 18 | 2893.0 | 30 |
| | | | | | | | | 4253.9 | 17 | 2992.7 | 29 |
| | | | | | | | | 4519.8 | 16 | 3099.6 | 28 |
| | | | | | | | | 4821.0 | 15 | 3214.3 | 27 |
| | | | | | | | | 5165.3 | 14 | 3337.9 | 26 |
| | | | | | | | | 5562.5 | 13 | 3471.4 | 25 |
| | | | | | | | | | | 3616.0 | 24 |
| | | | | | | | | | | 3773.2 | 23 |
| | | | | | | | | | | 3944.6 | 22 |
| | | | | | | | | | | 4132.4 | 21 |
| | | | | | | | | | | 4339.0 | 20 |
| | | | | | | | | | | 4567.3 | 19 |
| | | | | | | | | | | 4821.0 | 18 |
| | | | | | | | | | | 5104.5 | 17 |

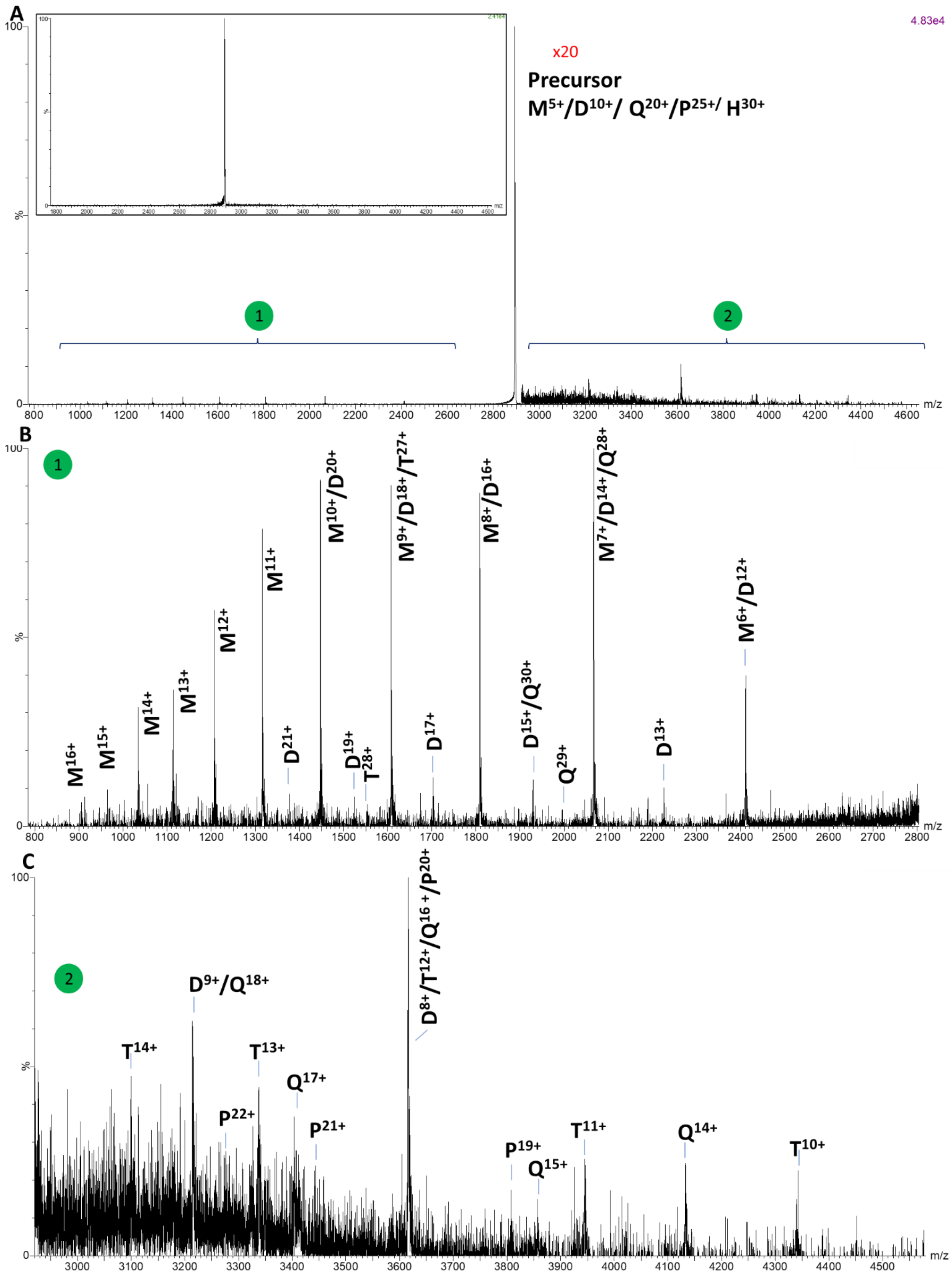


**Figure S6.** A, Native top-down CID fragmentation mass spectrum of m/z 2893 ion isolated in the quadrupole (precursor shown in inset on the left) and fragmented with 35 V in the trap region. The higher m/z region [2] of product ions is shown with x20 magnification here. B, expanded region of mass spectrum of product ions with lower m/z [1] and C, expanded mass spectrum of product ions with higher m/z [2]. The precursor $M^{5+}/D^{10+}/Q^{20+}/P^{25+}/H^{30+}$ dissociates giving rise to monomers and dimeric to pentameric oligomer populations with characteristic charge reduction. Data were acquired using 67 μM α-syn in 50 mM $NH_4OAc$, pH 7. M: monomer, D: dimer, T: trimer, Q: tetramer, P: pentamer and H: hexamer.

Collision induced dissociation mass spectra of the m the m/z 2893 ions under native-like conditions are shown in **Figure S6** revealed oligomeric populations up to hexameric units that dissociated with the release of monomers and charge reduced multimeric subunits during fragmentation in the trap region of the mass spectrometer.

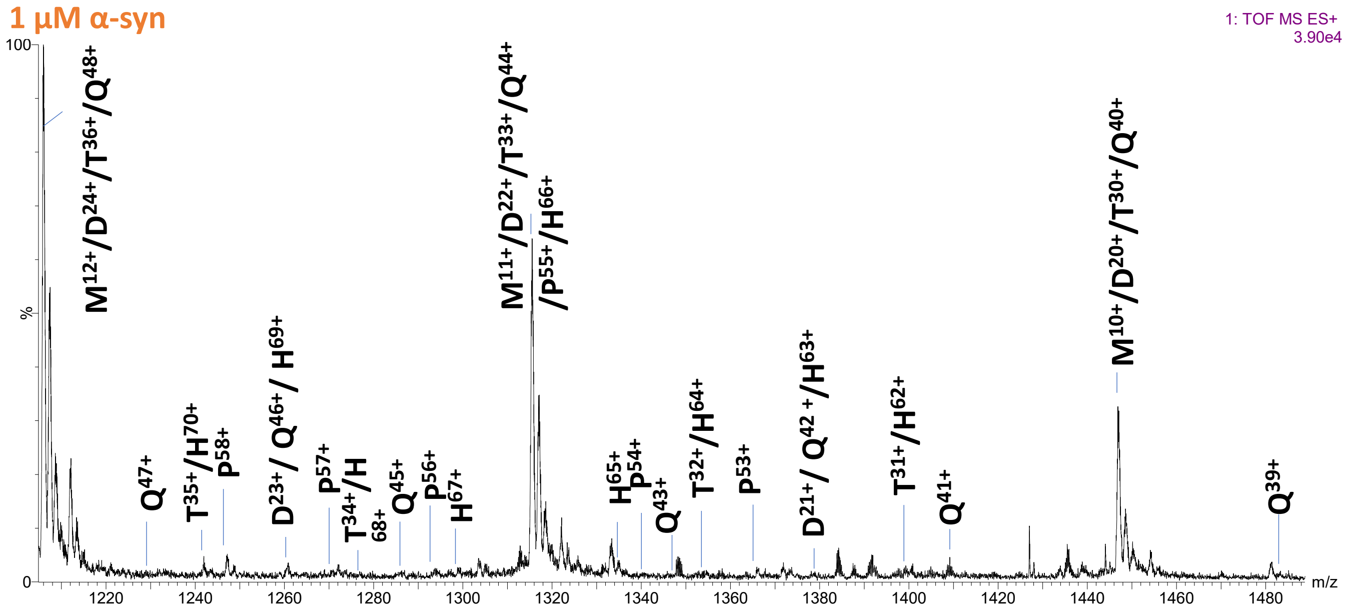


**Figure S7**. Expanded mass spectrum of 1 µM α-syn in 100 mM $NH_4OAc$, pH 7 under native-like conditions. Monomeric and oligomeric charge states are denoted with the following abbreviations: M: monomer, D: dimer, T: trimer, Q: tetramer, P: pentamer and H: hexamer.

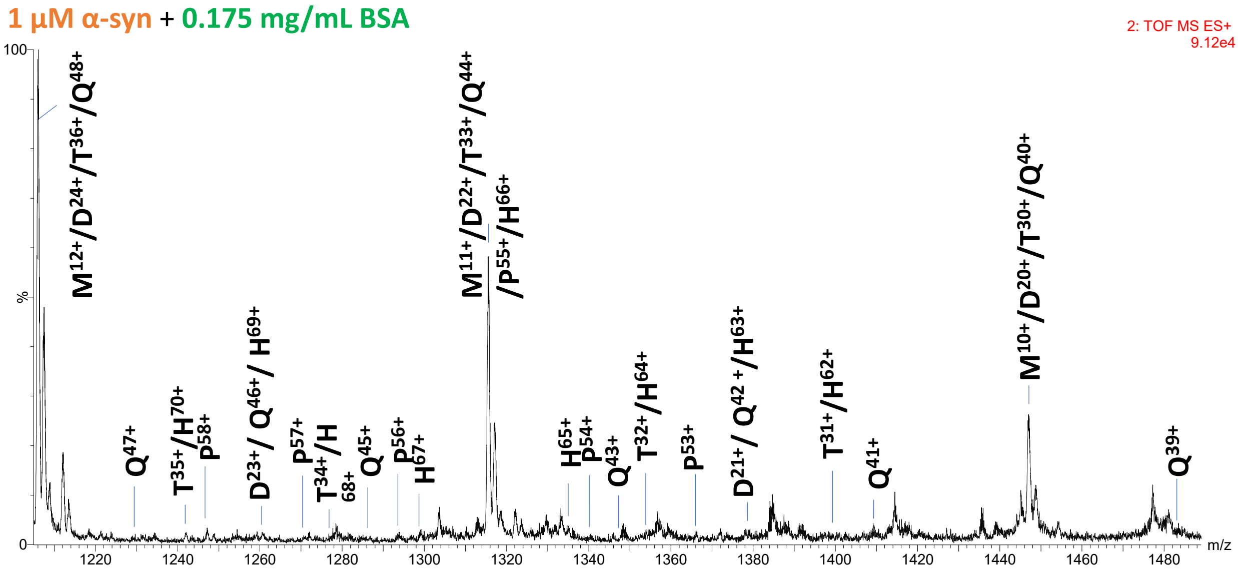


**Figure S8**. Expanded mass spectrum of 1 µM α-syn with 0.175 mg/mL BSA (2.6 µM) in 100 mM ammonium acetate, pH 7.0 under native-like conditions. Monomeric and oligomeric charge states are denoted with the following abbreviations: M: monomer, D: dimer, T: trimer, Q: tetramer, P: pentamer and H: hexamer.

Deconvolution of the intact mass also provides evidence for α-syn oligomers (**Figure S8 and Table S4**) at 67 µM concentration. To further ascertain the multimeric complexes of α-syn, collision induced dissociation mass spectra of the m the m/z 2893 ions under native-like conditions are shown in **Figure S5** revealing overlapping populations that dissociate with the release of monomers and charge reduced multimeric subunits during fragmentation in the trap region of the mass spectrometer.

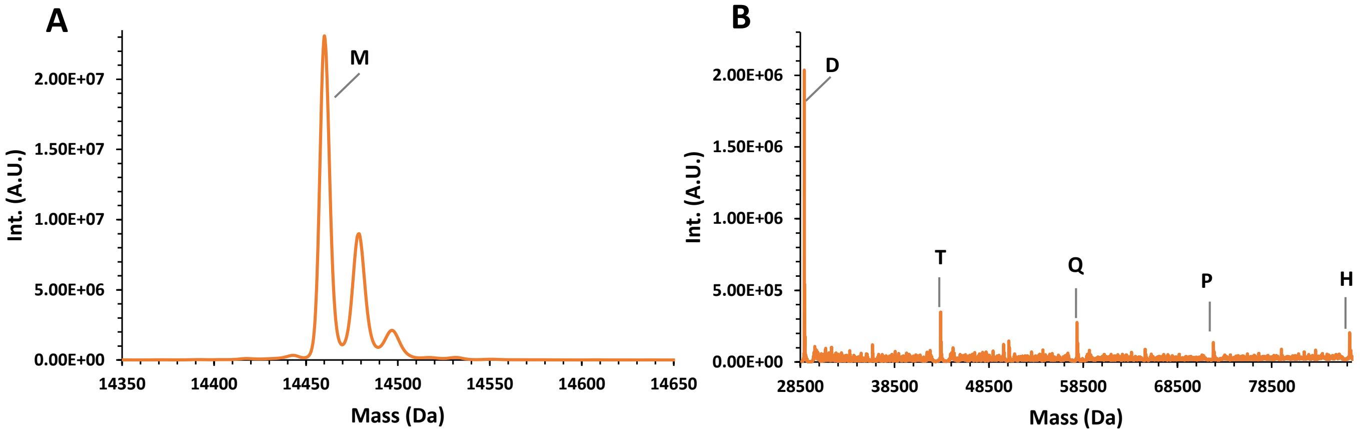


**Figure S9**. A, Deconvoluted mass of monomeric 67 µM α-syn in 50 mM $NH_4OAc$, pH 7 under native-like conditions. B, Deconvoluted mass spectrum of monomeric 67 µM α-syn in 50 mM $NH_4OAc$. Monomeric and oligomeric charge states are denoted as: M: monomer, D: dimer, T: trimer, Q: tetramer, P: pentamer and H: hexamer.

**Table S4**. Corresponding deconvoluted experimental masses of 67 µM α-syn in 50 mM NH4OAc, theoretical values and delta masses (Da).

| ID | deconv. mass | theo. mass | delta mass |
|---|---|---|---|
| monomer (M) | 14460 | 14460 | 0 |
| monomer (M) + 1 ammonia | 14479 | 14478 | 1 |
| monomer (M) + 2 ammonia | 14497 | 14496 | 1 |
| dimer (D) | 28920 | 28920 | 0 |
| trimer (T) | 43381 | 43380 | 1 |
| tetramer (Q) | 57840 | 57840 | 0 |
| pentamer (P) | 72300 | 72300 | 0 |
| hexamer (H) | 86760 | 86760 | 0 |

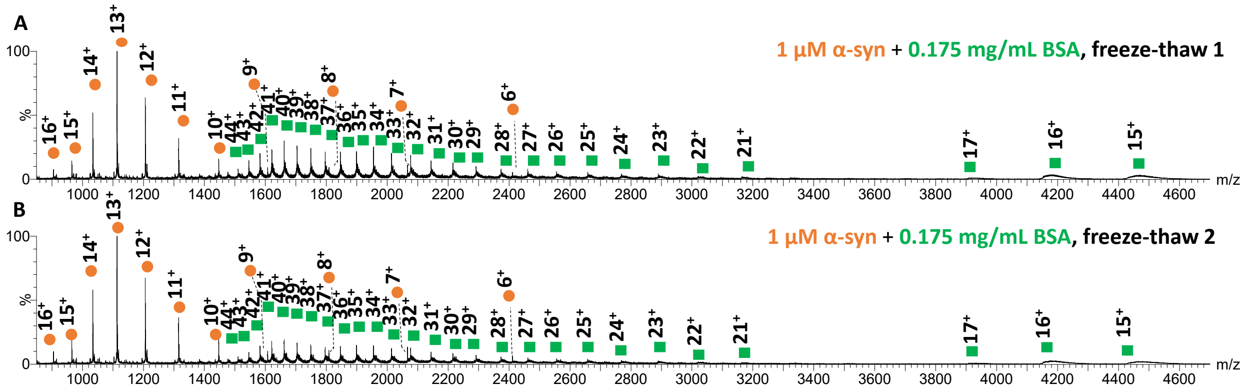


**Figure S10.** A: Mass spectrum of 1 µM α-syn with 0.175 mg/mL (2.6 µM) BSA in 100 mM ammonium acetate, pH 7.0 acquired following overnight storage at -80˚C and a thaw at room temperature. B: Mass spectrum of 1 µM α-syn with 0.175 mg/mL (2.6 µM) BSA in 100 mM ammonium acetate, pH 7.0 acquired following two cycles of freeze (-80˚C) and a thaw (at room temperature). All mass spectra shown were obtained at 38 V IMS wave height. α-syn charge states are shown with orange circles, bovine serum albumin monomers with green squares.

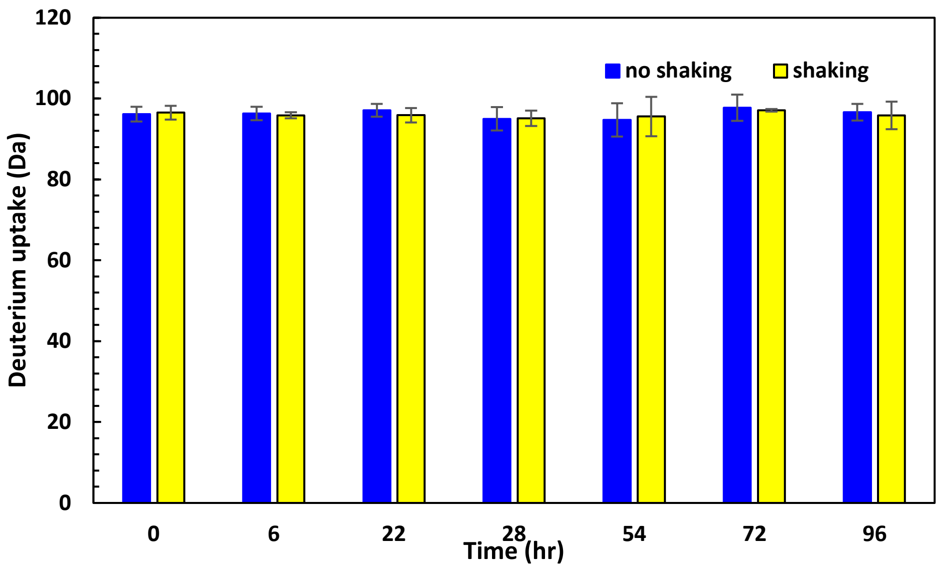


**Figure S11.** Mean deuterium uptake ± three standard deviations of α-syn in the presence of bovine serum albumin with or without shaking at 350 rpm on an orbital shaker at 21-22 ˚C. No statistically significant differences were observed between the two conditions following multiple t-tests of means (unpaired t-test of means) at the different timepoints (alpha was set at <0.05).

Previous studies describing aggregation of α-syn were conducted at much higher protein concentrations(9,10) and in the presence of vigorous agitation with a glass beads(9). Under our conditions, we did not observe a loss in deuterium uptake featuring a decreased solvent accessibility and hydrogen bonding.

**Table S5**. Skyline MRM optimisation. Optimal transitions for each peptide obtained using Skyline. Collision energies are in brackets.

| | Natural | | | Labelled | | |
|---|---|---|---|---|---|---|
| | MRM1 | MRM2 | MRM3 | MRM1 | MRM2 | MRM3 |
| T6 | 415.72>546.29 (17 eV) | 415.72>475.25 (16 eV) | 415.72>346.21 (16 eV) | 419.73>554.30 (17 eV) | 419.73>483.27 (16 eV) | 419.73>354.22 (16 eV) |
| T8 | 476.26>553.30 (15 eV) | 476.26>666.38 (15 eV) | 476.26>291.17 (15 eV) | 480.27>561.31 (15 eV) | 480.27>674.40 (15 eV) | 480.27>299.18 (15 eV) |
| T12 | 643.35>773.45 (20 eV) | 643.35>874.50 (20 eV) | 643.35>617.36 (20 eV) | 646.03>781.47 (20 eV) | 646.03>882.51 (20 eV) | 646.03>625.38 (20 eV) |
| T13 | 493.60>622.36 (14 eV) | 493.60>693.39 (14 eV) | 739.90>764.43 (26 eV) | 496.27>630.37 (14 eV) | 496.27>701.41 (14 eV) | 743.90>772.44 (26 eV) |

**Table S6.** Isobaric interferences in the CSF samples. MRM ratios of the natural channels of each CSF sample was compared with the calibration standard.

T12

| | MRM2/MRM1 Ratio | MRM3/MRM1 Ratio | MRM2/MRM3 Ratio |
|---|---|---|---|
| NeuroMet003 | 0.89 | 0.59 | 1.50 |
| NeuroMet018 | 0.92 | 0.69* | 1.36 |
| NeuroMet019 | 0.88 | 0.57 | 1.56 |
| NeuroMet021 | 0.92 | 0.59 | 1.56 |
| NeuroMet024 | 0.90 | 0.60 | 1.51 |
| NeuroMet025 | 0.93 | 0.59 | 1.57 |
| NeuroMet030 | 0.92 | 0.56 | 1.64 |
| NeuroMet035 | 0.95 | 0.57 | 1.68 |
| NeuroMet042 | 0.86 | 0.58 | 1.49 |
| NeuroMet045 | 0.92 | 0.57 | 1.60 |
| NeuroMet046 | 0.89 | 0.55 | 1.61 |
| NeuroMet074 | 0.90 | 0.59 | 1.54 |
| NeuroMet080 | 0.94 | 0.56 | 1.68 |
| NeuroMet085 | 0.94 | 0.58 | 1.64 |
| NeuroMet087 | 0.90 | 0.55 | 1.63 |
| STD AVG | 0.92 | 0.57 | 1.60 |

T13

| | MRM2/MRM1 Ratio | MRM3/MRM1 Ratio | MRM2/MRM3 Ratio |
|---|---|---|---|
| NeuroMet003 | 0.77* | 1.01 | 0.76 |
| NeuroMet018 | 0.88 | 1.04 | 0.85 |
| NeuroMet019 | 0.99 | 1.21 | 0.82 |
| NeuroMet021 | 1.10 | 1.23 | 0.90 |
| NeuroMet024 | 0.96 | 1.16 | 0.83 |
| NeuroMet025 | 0.95 | 1.13 | 0.84 |
| NeuroMet030 | 1.00 | 1.14 | 0.87 |
| NeuroMet035 | 0.94 | 1.16 | 0.81 |
| NeuroMet042 | 0.95 | 1.14 | 0.82 |
| NeuroMet045 | 1.01 | 1.28 | 0.79 |
| NeuroMet046 | 1.02 | 1.23 | 0.83 |
| NeuroMet074 | 1.04 | 1.24 | 0.84 |
| NeuroMet080 | 0.95 | 1.21 | 0.78 |
| NeuroMet085 | 0.96 | 1.23 | 0.78 |
| NeuroMet087 | 0.93 | 1.15 | 0.81 |
| STD AVG | 1.00 | 1.11 | 0.90 |

STD AVG = average ratios from the upper portion of the calibration standard (2 ng/g to 10 ng/g)
* = >15% deviation from the standard average

**Table S7.** Recovery values (%) of four α-syn synuclein peptides.

| Peptide | Recovery |
|---|---|
| T6 | 77% |
| T8 | 75% |
| T12 | 73% |
| T13 | 67% |

**S12 Materials and Data Processing**

Chemical and reagents

The α-syn potential calibrator was commercially available from Promise Advanced Proteomics (Grenoble, France). The isotopically labelled α-syn (13C and 15N) was also purchased from Promise Advanced Proteomics (Grenoble, France). The purity of the primary calibrator confirmed using high resolution mass spectrometry by using both a Waters H-Class ultra-performance liquid chromatographic (UPLC) system coupled to a Waters Xevo G2-XS QToF mass spectrometer (Waters, Wilmslow, UK) operated in a sensitivity mode with a resolving power of 25,000 or a Thermo Fisher Scientific Vanquish binary UPLC system coupled to a Thermo Fisher Scientific Q-Exactive Plus (Thermo Fisher Scientific, Bremen, Germany) operated at 30,000 resolution mode. All analyses were performed in full scan MS mode and impurities identified based on accurate mass. Other materials were: ultrapure water (18 mol/L Ω $cm^{-1}$), formic acid (Fisher Scientific methanol, acetonitrile (Honeywell). All solvents were LC-MS grade. Reagents for artificial CSF were: sodium chloride, sodium bicarbonate, potassium chloride, sodium phosphate monobasic, magnesium chloride, D-(+)-Glucose and calcium chloride (Sigma-Aldrich).

Cerebrospinal fluids (CSF)

A pool of 100 CSF samples from the biological resource centre (CRB) of CHU-Montpellier was used for validation experiments. Samples for generating pools were retrospectively selected from the NeuroCognition Biobank of Montpellier's University Hospital from patients recruited in the Resource and Research Memory Centre. Patients gave informed and written consent to have their samples stored in an officially registered and ethically approved biological collection (#DC-2008-417) and later used for scientific research. A panel of fifteen CSF samples were obtained from Charité which were analysed using the candidate reference method and the U-PLEX Human α-Synuclein Kit (Mesoscale) following the manufacturer's instructions. The antibodies used in the kit are directed against the C-terminal part of the α-syn (110 – 125 aa) for the rabbit monoclonal capture antibody and the mouse monoclonal detection antibody captures the residues between 15 – 125 aa.

**Data acquisition and processing**

IMS-MS

Myoglobin from equine skeletal muscle (M0630), ubiquitin from bovine erythrocytes (U6253), and cytochrome c from equine heart (C2506), bovine serum albumin (B4287) were purchased from Sigma-Aldrich (now Merck, Gillingham, UK) and ammonium acetate (A/3440/50) from Fisher Scientific (Loughborough, UK). Recombinant α-syn was provided by UCL as a gift.

nESI sampling was performed by use of the Triversa Nanomate (Advion, Ithaca, NY, USA) platform interfaced with a Synapt G2Si mass spectrometer (Waters Corp, Wilmslow, UK), via an electroconductive tip coupled to a chip by direct infusion from a 96 well microtiter plate. The Triversa NanoMate robot was operated with the ChipSoft 8.3.3 software at 1.65 kV capillary voltage and 0.35 psi pressure. Protein stocks were prepared by diluting the 1 mg/mL α-syn stock stored frozen in water at -80 ˚C and adjusting its concentration based on its molar extinction coefficient (5960 $M^{-1}$ $cm^{-1}$) measured on a Nanodrop 2000 spectrophotometer at 280 nm wavelength. Denatured samples of protein standards (ubiquitin, myoglobin and cytochrome c) were prepared with 50% acetonitrile, 49% water, and 1% formic acid, under acidic conditions at pH 2.2. Native-like samples were prepared using either 50 mM or 100 mM ammonium acetate ($NH_4OAc$), pH 7.0 in ultrapure water (18 MΩ $cm^{-1}$) (Elga Systems, Lane End, UK). A 5 mg/mL bovine serum albumin stock solution was prepared in water and diluted at final concentration of 0.175 mg/mL in 100 mM ammonium acetate. The protein solutions were analysed immediately after preparation, or in the case of the freeze and thaw samples the solution was kept at -80 ˚C for one period of overnight storage or two followed by analyses. The Synapt G2Si (Waters Corp., Wilmslow, UK) mass spectrometer was operated in positive electrospray ionisation mode, at a helium flow rate of 180 mL/min and nitrogen flow rate of 90 mL/min. External mass calibration was performed with caesium iodide in the corresponding mass ranges. All measurements were conducted in triplicate, accumulating 90−100 scans with a 1 s scan time. CCS calibration was performed with ubiquitin, myoglobin and cytochrome c following the protocol described by Ruotolo et al.(11). CCS are reported as ${}^{TW}CCS_{N2\rightarrow He}$, that is, helium reference values have been used to calibrate TWIMS measurements made in nitrogen. Reference CCS values were obtained from the Bush database (myoglobin) and the Clemmer database (ubiquitin and cytochrome c). Mass spectra and drift times were not smoothed unless specified. The following settings were used: capillary voltage 1.80 kV, cone 70 V, source temperature 100°C, trap collision energy 4 V, transfer collision energy 2 V, trap wave velocity 311 m/s, wave height 6 V, IMS wave velocity 650 m/s, wave height 38-40 V, transfer wave velocity 190 m/s, wave height 4 V, trap bias 45, IMS bias 3, step wave 1 velocity 300 m/s, step wave 1 height 10 V, step wave 2 velocity 300 m/s, step wave 2 wave height 0 V, step wave 1 RF offset 300, step wave 2 RF offset 380, backing pressure 2.82 mbar and m/z range 200-6500. All data were acquired in triplicate and in MassLynx V.4.1 (Waters, Wilmslow, UK).

Collision induced unfolding (CIU)

The precursor ions (10+ and 12+) were selected manually in the quadrupole in MS/MS mode. All mass spectrometry conditions were the same as described above for IM-MS section with the exception of applying MS/MS mode. Collision induced unfolding of 1 µM α-syn in 100 mM ammonium acetate and 1 µM α-syn in 100 mM ammonium acetate with 2.6 µM BSA was initiated by ramping the voltages 4 V – 26 V in the trap region of the instrument by manual increments of 2 V in a randomized order. All experiments were repeated three times. 80 scans were acquired at each voltage at 1 s scan rate. Data were analysed in CIU Suite 2. 2 with a smoothing window of 5, iteration 2, and axis scaling step of 2 of collision voltage for interpolation.

HDX-MS

Materials: Sodium phosphate monobasic (205925000), sodium phosphate dibasic (215472500) were obtained from Acros Organics (Geel, Antwerp, Belgium), deuterium oxide ($D_2O$) 99.9% (450510), Glu-fibrinopeptide B, human (F3261) and urea (U5378) were purchased from Sigma-Aldrich (now Merck, Gillingham, UK); 1.5 mL LoBind Eppendorf tubes (022431081) from Fisher Scientific (Loughborough, UK), deuterium chloride (175420500) for pH adjustment was purchased from Acros Organics (Geel, Antwerp, Belgium) and hydrochloric acid (318965) from Fluka Analytical (Honeywell Research Chemicals, Seelze, Germany). Maximum recovery vials (VI-04-12-06 MRL) were purchased from Chromatography Direct (Runcorn, UK) and amber crimp vials (10003264) from Fisher Scientific (Loughborough, UK). Recombinant α-syn was provided by UCL as a gift.

45 µL 15 µM protein was incubated at pH 7.0 at 21-22 °C in LoBind Eppendorf tubes (022431081) (Fisher Scientific, Loughborough, UK) in the presence of 0.175 mg/mL bovine serum albumin (2.6 µM) with or without shaking at 350 rpm on an orbital benchtop shaker (Grant-Bio PMS 1000i, Shepreth, UK). The fibrillation process or lack of thereof was monitored by taking temporally resolved samples of α-syn (timepoints: 0 h, 6 h, 22 h, 28 h, 54 h, 72 h and 96 h) and snap-frozen for HDX-MS. The samples were thawed at room temperature prior to mass spectrometry. To initiate deuterium labelling, 12 µL of the above sample, one at a time, was transferred into a maximum recovery vial (VI-04-12-06 MRL) (Chromatography Direct, Runcorn, UK), placed into a LEAP PAL

robot (LEAP Technologies, Morrisville, North Carolina, USA) and diluted into 135 µL of 10 mM potassium phosphate buffer composed of potassium phosphate monobasic and dibasic salts made in $D_2O$, pH 7.0, and incubated at 21-22 °C for 1 min. After a one-min pulse, 50 µL of the above sample was mixed with 50 µL of chilled quench buffer comprising of 6 M urea with 5% formic acid prepared in 100 mM potassium phosphate buffered, consisting of potassium phosphate monobasic and dibasic salts, in amber crimp vials (10003264) (Fisher Scientific, Loughborough, UK). 95 µL of quenched sample was injected onto a temperature-controlled nanoACQUITY UPLC System with HDX technology (Waters, Milford, Massachusetts, USA) chromatographic separation. Sample handling and mixing steps were performed using a first-generation LEAP PAL system set up for HDX analysis. The samples were injected, loaded at 50 µL/min and trapped at 80 µL/min for 3 mins in ultrapure water with 0.5% (v/v) formic acid and subsequently eluted using a linear gradient on a MassPREP microdesalting column (186004032) (Waters, Wilmslow, UK). The flow rate was 100 µL/min (acetonitrile with 0.5% (v/v) formic acid). The column was held at 0 °C. Chromatographic separation was performed at 100 µL/min using an 8 min gradient from 80% A/20% B to 10% A/90% B. The source conditions were: temperature 100 ˚C, desolvation temperature 250 ˚C, cone gas flow 99 L/min, desolvation gas flow 499 L/min and trap gas flow 2.4 mL/min. Data were acquired in the presence of a lockmass (GluFib) in the range of 100 – 2500 m/z, in resolution mode. Data were imported into DynamX v3.0 (Waters, Milford, Massachusetts, USA) to generate uptake plots. The shift in the mass of labelled peptide relative to the unlabelled peptide was used to determine the extent of deuterium incorporation. Prolines and the first two residues were excluded from exchange, giving 134 exchangeable sites for α-syn.

Deconvolution

For deconvolution the embedded MaxEnt1 function of MassLynx v4.1 (Waters Ltd, Wilmslow, UK) was used in the mass range of 10800-89300 Da, with a resolution channel of 1 Da, using the damage model with a uniform Gaussian peak width at half height of 0.750 Da, minimum intensity ratios were set on the left at 10% and on the right at 40%, performing 10 iterations.

Native top-down mass spectrometry

Precursor ion m/z 2893 of 67 µM α-syn in 50 mM $NH_4OAc$, pH 7.0 was isolated in the quadrupole region of a Synapt G2 Si mass spectrometer (Waters Ltd, Wilmslow, UK) operated in positive mode and fragmented in the trap region at 28 V and 35 V respectively. The precursor ions were collected for 1.4 min and 1.8 min respectively, and fragment ions for 10-11 mins. The settings were: Capillary 1.80 KV, cone 50 V, sampling temp. 100 ˚C, offset 70, desolvation temperature 250 ˚C, cone gas flow 150 L/h, purge gas flow 500 mL/h, desolvation gas flow 500 L/h, nebuliser gas flow 7 bar, trap 4 V, transfer 2 V for precursor, trap DC bias 45, trap wave velocity 311 m/s, trap wave height 6 V, IMS wave velocity 650 m/s, IMS wave height 40 V, step wave 1 velocity 300 m/s, step wave 1 height 10 V, step wave 2 velocity 300 m/s, step wave 2 height 0 V, step wave 1 RF offset 300, step wave 2 RF offset 380, backing 3.17 mbar, m/z 600- 5500, 1 s scan time, helium flow rate 180 mL/min and nitrogen flow rate 90 mL/min. Mass spectra were viewed and processed in in MassLynx V.4.1 (Waters Corp, Wilmslow, UK) with the application of smoothing (smooth window size (scan) ±5, number of smooths 1, Savitzky Golay algorithm). External mass calibration was performed with caesium iodide in the corresponding mass ranges.

Quantitative nuclear magnetic resonance (qNMR)

Approximately 2 mg of peptide was weighed into an aluminium boat and transferred to a LoBind Eppendorf tube. An aliquot (0.2 mL) of a maleic acid internal standard solution was accurately weighed in to the LoBind tube and approximately 0.3 mL of $D_2O/CD_3CN$ was added. The resultant solution was vortexed and transferred to a 5 mm NMR tube for analysis. Two independent solutions were prepared for T6. Considering the acquisition time for the qNMR experiments (~ 45 min each) and unknown stability of the solutions, two qNMR experiments were acquired for the replicate 1 solution, and a single qNMR experiment was acquired for the replicate 2 solution. The $^{1}$H qNMR analyses were performed on a Bruker Avance 600 MHz NMR. The following experimental parameters were used:

| | |
|---|---|
| Number of scans: | 64 |
| Relaxation delay: | 40 seconds |
| Spectral width: | 20.6 ppm |
| Temperature: | 298.0 K |
| FID processing software: | Topspin 3.5 pl 2 |
| Baseline correction: | Manual, polynomial baseline correction |
| Signal integration: | Manual |

The uncertainty contribution of the qNMR quantification was calculated to be 1%.

Optimisation of MRM transitions using Skyline

aCSF (500 µL, 0.175 mg/mL BSA), was spiked with primary calibrator to a final concentration of 1.5 ng/g and labelled α-syn (100 µL, 3.3 nM). The sample digested with trypsin (3.5 µg) overnight at 27 °C. The peptides cleaned up using SPE, eluted into two fractions (T6/T12/T13 and T8) and dried to completion. The full amino acid sequence of α-syn was imported into Skyline and the relevant peptides were highlighted. The default transitions (y-ions) of the monitored peptides were isolated and further y-ions were added to a maximum of 90 MRMs at any time. The previously prepared samples were ran using the newly created MS method and the top three transitions for each peptide were taken further for collision energy optimisation.

Measurement Uncertainty

The standard and combined uncertainties were calculated in accordance with the EURACHEM/CITAC Guide CG4 and with the ISO Guide to the Expression of Uncertainty in Measurements.(12,13)

The uncertainty (u) associated with the double exact matching isotope dilution mass spectrometry values from the amino acid analysis or tryptic digestion experiments for quantification of the α-syn primary calibrator were calculated as previously reported.(14,15) Particularly the combined uncertainty associated to the α-syn primary calibrator was calculated as by equation 1:

$$u = \sqrt{u_{av}^2 + b_{var}^2}$$

$u_{av}$ is the average standard uncertainty calculated from nine tryptic digests for the three peptides monitored (T1, T6 and T8) and $b_{var}$ represents the variability of the digestion and is calculated by equation 2:

$$b_{var} = \sigma\sqrt{n}$$

$\sigma$ is the standard deviation from the α synuclein values obtained by n tryptic digestions from each peptide (T1, T6 and T8).

The uncertainty from the calibration curve measurements was estimated based on the Eurachem guide to Quantifying Uncertainty in Analytical Measurement.(14) Briefly the standard deviation of the calibration curve residuals was calculated using equation 3:

$$s = \sqrt{\frac{\sum_i (y_i - \hat{y}_i)^2}{n - 2}}$$

$y_i$ is the peak area ratio between the natural and labelled peptides, $\hat{y}_i$ is the value of y generated from the calibration curve at a gravimetrically calculated primary calibrator concentration and $n$ is the number of calibration measurements.

The uncertainty associated with the calibration curve quantification was estimated for each of the three peptides quantified using Equation 4:

$$u(c_0) = \frac{s}{b}\sqrt{\frac{1}{m} + \frac{1}{n} + \frac{(y_0 - \bar{y})^2}{b^2 \sum_i (x_i - \bar{x})^2}}$$

$s$ is the residual standard deviation of the calibration curve obtained from equation 3, $b$ is the slope of the calibration curve, $m$ is defined as the number of analytical measurements performed on the sample and $n$ is the number of calibration measurements. $x_i$ is the measured α-syn sample concentration, $\bar{x}$ is the average concentration of all the calibration standards, $y_0$ is the observed peak area ratio between the natural and labelled peptides and $\bar{y}$ is the average natural to labelled ratio of the calibration curve.

Reference method validation (Recovery)

Recovery of the monitored peptides was estimated by spiking labelled α-syn pre- and post-clean-up. Three conditions were prepared in triplicate in 500 µL aCSF (0.175 mg/mL BSA), including a pre-SPE sample, post-SPE sample and control sample. The pre- and post-SPE samples were only spiked with natural protein (1.5 ng/g), while the control was spiked with both natural and labelled (both at 1.5 ng/g). In a separate vial, labelled protein (1.5 ng/g) was prepared in buffer (50 mM AMBIC). Finally, a blank vial with only buffer (50 mM AMBIC) was prepared. The pre-SPE, post-SPE and control samples were basified using tris buffer (1 M, pH 8.2) to a final Tris concentration of 50 mM. The natural and labelled samples were digested using trypsin (Promega) at a 1:25 trypsin to protein ratio overnight at 27 °C. The pre-SPE samples were spiked with labelled digests, whereas the post-SPE samples were spiked with a blank digest prior to the clean-up. After the SPE process, labelled digests were added to the post-SPE samples, whereas the other set of samples was spiked with an equivalent blank digest. Each sample was injected in triplicates. The MS ion signal area ratio between the natural and labelled α-syn was corrected for the gravimetric amount of labelled α-syn added. The corrected ratios were averaged for the pre- and post-SPE sets of samples to get two values which were then used to calculate the recovery. The pre-SPE ratio was divided by the post-SPE ratio, to give a percentage value that represents recovery. The results for the four monitored peptides are shown in the following table.
